# Accurate Probabilistic Reconstruction of Cell Lineage Trees from SNVs and CNAs with ScisTreeCNA

**DOI:** 10.1101/2025.11.21.689819

**Authors:** Haotian Zhang, Yufeng Wu

## Abstract

Cell lineage tree is a fundamental evolutionary model for single-cell evolution. Inference of cell lineage tree from noisy single-cell DNA data has been studied actively in recent years. Existing methods for cell lineage tree inference can be classified into two categories based on the type of genetic variations they work with: single-nucleotide variants (SNVs) or copy-number aberrations (CNAs). Due to various noises and uncertainties in the data, the existing methods are not fully satisfactory, in part because they only used one type of genetic variant. Single-cell DNA sequencing data with both SNVs and CNAs are becoming available. In principle, joint inference of cell lineage trees from both SNVs and CNAs may lead to more accurate results. However, there is a lack of rigorous models and efficient algorithms for such inference. In this paper, we present a new cell lineage tree inference method, called ScisTreeCNA, that jointly infers cell lineage trees from SNVs and CNAs. A key contribution of ScisTreeCNA is a novel probabilistic model for the joint evolution of SNVs and CNAs in single cell data. Based on this model, ScisTreeCNA implemented several efficient algorithms for accelerating probabilistic inference of cell lineage tree. Experiments on both simulated and real biological data show that ScisTreeCNA consistently outperforms existing methods in the accuracy of the inferred cell lineage trees. ScisTreeCNA is available at https://github.com/haotianzh/ScisTreeCNA.

## 1 Introduction

The fast-developing single-cell sequencing technologies are enabling novel applications in several important areas of biology, including cancer biology and developmental biology (e.g., [1, 2]). One of the most important applications is the study of evolution at the single-cell level. Suppose that a group of single cells has been sequenced using single-cell DNA sequencing (scDNA). The evolutionary history of these cells can be modeled by a rooted phylogenetic tree, called *cell lineage tree* (CLT). The leaves of a CLT are the extant sampled cells, while the internal nodes are the ancestral cells. CLT is a fundamental model in single-cell evolution, and has been used widely in areas such as cancer genetics [3].

While not directly observable, CLT can be inferred from the genetic variants called from scDNA data, including single nucleotide variations (SNVs) and copy number aberrations (CNAs). Single-cell data tend to be very noisy and sparse. Most existing phylogenetic inference methods are not directly applicable for noisy scDNA data. Therefore, inference of CLT from noisy data is currently a very active research subject.

Most existing CLT inference methods are designed to work with either SNVs or CNAs, but not both. There are many SNV-based methods, including SCITE [4], ScisTree [5], CellPhy [6] and ScisTree2 [7], among others (e.g. [8, 9]). For CNA data, the classic neighbor joining [10] is often used, along with several other methods (e.g., [11, 12]). While these methods are certainly useful, the accuracy of the inferred CLT may still be low for data with significant noise [11, 13]. In many scDNA-seq datasets, SNVs often co-occur with CNAs [13], meaning that CNA events may also affect SNV loci. Many SNV-based methods [4, 5, 7] assume the infinite-sites (IS) model, in which each SNV is acquired once and never lost during evolution. However, CNAs can cause violations of this assumption [13]. SNV-based methods that instead adopt the finite-sites (FS) model (e.g., [6]) are less affected by the presence of CNAs, but the FS model may be less phylogenetically informative than the IS model [13]. Other approaches, such as those based on the Dollo model (e.g., [14]), assume that an SNV can be gained once but lost multiple times. Nevertheless, the Dollo model still does not fully capture the effects of copy number changes.

There are scDNA-seq datasets that contain both SNVs and CNAs, such as those generated using Mission Bio technologies [15]. Another example is the high-grade serous ovarian cancer (HGSOC) dataset from [16], which includes both SNV and CNA profiles for over 800 cells.

A method that integrates both SNVs and CNAs may infer a more accurate CLT, as it leverages a richer set of genomic information than existing approaches. Currently, no practical CLT inference methods jointly model both SNVs and CNAs. The closest related approaches that incorporate both data types focus instead on reconstructing clonal trees in cancer genomics (e.g., [13, 17, 18, 19, 20]). For example, ConDoR [18] first clusters cells into clones using CNAs and then reconstructs clonal trees with SNVs. In principle, such approaches can be applied to reconstruct CLT. However, methods such as ConDoR are not computationally efficient for inferring CLTs that are usually much larger than clonal trees. ConDoR only considers copy losses but not gains. Moreover, to reconstruct a clonal tree, cells need to be assigned to clones first. But there is uncertainty in how to assign cells into clones.

In this paper, we introduce ScisTreeCNA, a novel maximum likelihood inference method for cell lineage tree inference jointly from SNVs and CNAs. On the high level, ScisTreeCNA retains the local search approach for optimization in our previous ScisTree2 approach [7]. ScisTreeCNA, however, is more sophisticated than ScisTree2 in methodology with the following features:

- A novel probabilistic model (named as the *JSC* model) for the joint evolution of SNVs and CNAs on a CLT.
- Efficient algorithms for maximum likelihood inference of CLT based on the JSC model.
- Further acceleration of the likelihood computation using GPUs.

Experiments on simulated data and two real biological datasets demonstrate that ScisTreeCNA achieves higher accuracy in inferring CLTs from noisy scDNA-seq data compared to existing methods. Moreover, ScisTreeCNA is efficient and can handle datasets comprising hundreds of cells and sites. To the best of our knowledge, it is the first probabilistic inference method that infers CLT jointly from both SNVs and CNAs and is applicable to current scDNA-seq data.

## 2 Model

### 2.1 Generalized genotypes

At an SNV site *s*, an allele of a cell *c* may be wild-type (0) or mutant (1). With CNAs, the numbers of 0 or 1 alleles can be any non-negative integers. We use an ordered pair of integers (*g*_0_, *g*_1_) to represent the generalized genotype at a site and a cell. Here, *g*_0_ (respectively *g*_1_) is the number of wild-type (respectively mutant) alleles. For example, the rightmost genotype (2, 1) in Fig. 1 (a) has two wild-type alleles and one mutant allele.

**Figure 1.**
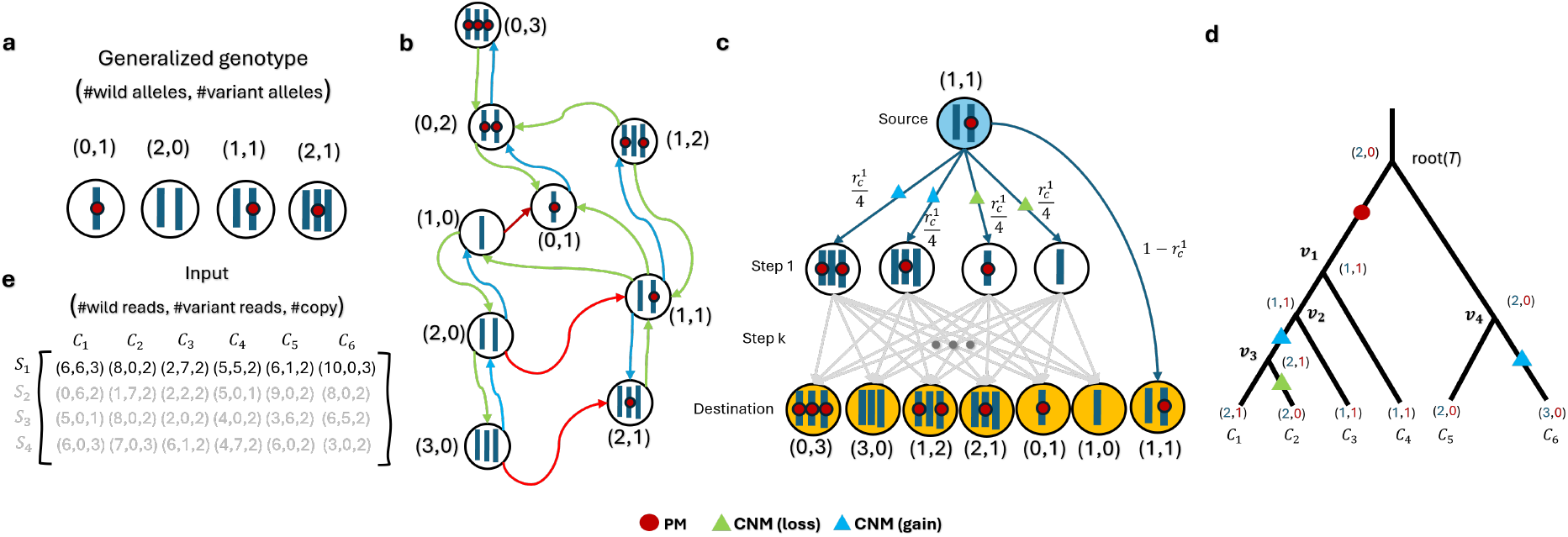
Generalized genotypes and mutations on cell lineage trees. (a) Generalized genotypes: pairs of integers as the numbers of wild-type and mutant alleles. (b) All possible genotypes with copy number within the range [1, 3] (i.e., p = 1 and q = 3) and how they transit by PM or CNM events. A PM changes a wild-type allele to a mutant allele. A CNM event increases or decreases the copy number by one. (c) Genotype transitions along a branch of CLT starting from genotype (1,1), with first-step transition probabilities annotated on the edges. Last row: final genotypes. (d) CLT of 6 cells annotated with PM, CNM, and the corresponding genotypes. (e) Input data of ScisTreeCNA for 6 cells and 4 sites: read counts of 0 and 1 alleles and the called copy numbers for each site and cell.

There are two types of mutations: point mutation (PM) and copy number mutations (CNM). A PM at *s* changes a 0 allele into a 1 allele. Here, we assume no back or recurrent PM at *s* by the IS assumption. We model CNM as the process of choosing a random allele and either duplicating (i.e., copy gain) or discarding (i.e., copy loss) the chosen allele. A genotype can transit to another by mutations. For example, in Fig. 1 part (b), the genotype (2, 0) can transit to (1, 1) by PM, to (1, 0) by CNM (loss) or to (3, 0) by CNM (gain). We denote the copy number of genotype (*g*_0_, *g*_1_) as *z*, where *z*(*g*_0_, *g*_1_) = *g*_0_ + *g*_1_. We assume that copy numbers are within a range [*p, q*] where *p* and *q* are known. For example, in Fig. 1 (b), *p* = 1 and *q* = 3. The copy number of the genotype (1, 2) is 3.

### 2.2 The JSC model

#### High-level idea

The JSC model is a generative model and specifies how stochastic mutational (PM and CNM) processes determine genotypes at each node in *T* . Starting from the root of *T* in a top-down manner, the genotype at each (leaf or internal) node is determined by where the single PM and potentially multiple CNMs occur in *T* . The genotypes at leaves can be estimated from their read counts and called copy numbers. While probabilistic models at single nucleotide level are commonly used [21], there are much less work on probabilistic models for copy number variations/aberrations. Existing models for copy numbers mainly focus on breakpoints (see, e.g., [22]). To the best of our knowledge, the JSC model is the first joint probabilistic model for the evolution of SNVs and CNAs.

We now define the JSC model and its parameters. We have the following assumptions which are commonly made in the phylogenetics literature (e.g., see [21]). For a fixed site,

1. A point mutation (PM) that changes a wild-type allele (0) to a mutant allele (1) can only occur once (the IS assumption). That is, the mutation can only be *gained once*.
2. A copy number mutation (CNM) takes an arbitrary (wild-type or mutant) allele and with *equal* probability, either (i) deletes this allele, or (ii) makes a copy of this allele.
3. All mutations (PMs or CNMs) are treated as *independent* because the sites in current data are often far apart.
4. Any mutational (PM or CNM) process along a *single* branch of *T* follows the standard *Poisson process*. More than one mutation may occur along a branch.
5. We assume that the genotype at the root of *T* is diploid.

#### 2.2.1 Poisson processes for the joint SNV and CNA evolution along a branch of *T*

We focus on a single SNV site *s* and a branch *b* = (*u, v*) in *T, u* is the source of *b*, and *v* is the destination of *b*. We consider the mutational (PM and CNM) events *forward in time*. The *waiting time* of the occurrence of a PM or CNM follows an *exponential* distribution due to the Poisson assumption (see, e.g., [23]). We let the arrival rates of PM (respectively CNM) be *λ*_*s*_ (respectively *λ*_*c*_).

##### Branch length modeling

When *b*’s length is infinite, a mutation event will occur eventually. However, *b*’s length is usually finite. Directly incorporating branch length into mutational processes complicates probabilistic modeling. Our idea for modeling the effect of finite branch length is introducing a *third* stochastic event: *timeout*, whose arrival time is modeled by an *exponential* distribution with rate *λ*_*t*_. The timeout is an *absorbing* state: once a timeout occurs, no more mutation events are allowed. Here, *λ*_*t*_ is influenced by the length of *b*. In *T*, the lengths of different branches can vary. To allow fast inference on this model, we assume in this paper that *λ*_*t*_ corresponds to the average branch length and remains the same for each *b*, although it is in principle straightforward to vary *λ*_*t*_ for different branches.

Along *b*, there is a sequence of *k* stochastic events, each of one of the three types: PM, CNM and timeout. Here, *k ≥* 1 and the last event is the timeout event. The first *k* − 1 events are either PM or CNM events. Now, suppose we are at a specific point *t*_*b*_ along *b*. We denote *E*_*M*_ as the type of the first coming event forward in time from *t*_*b*_. There are two different cases on the allowed types of *E*_*M*_ . It is well known in probability theory that the probability of the first event being of a certain type (say CNM) is *proportional* to the rate of the corresponding process when arrival of events follows the *exponential* distribution (see, e.g., [23], Sect. 2.2).

##### Case (i): CNM and timeout only

Sometimes PM is not allowed at *t*_*b*_ due to the IS model. This can be the case when the single PM has already occurred along *another* branch in *T* or *prior* to *t*_*b*_ along *b*. Then,

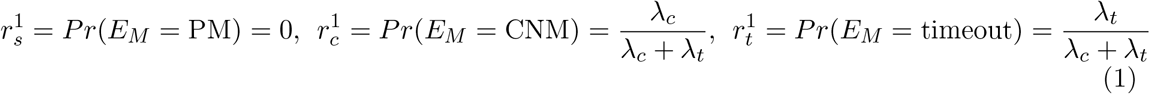

##### Case (ii): PM and CNM

It is also possible that only PM and CNM are allowed after *t*_*b*_. This occurs when the PM is known to occur after *t*_*b*_. For example, suppose that there is no 1 allele at the source *u* but there is a 1 allele at the destination *v*. Then a mutation must have occurred along *b*. Note that in this case timeout cannot occur until the PM occurs. Then,

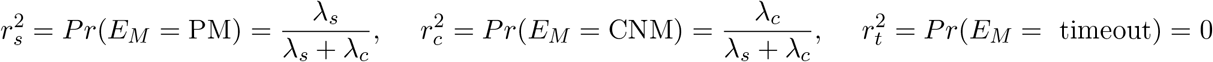

For example, in Fig. 1 (d), case (i) applies for (*v*_1_, *v*_2_) and case (ii) applies for (*root*(*T*), *v*_1_).

#### 2.2.2 Transition probability of genotypes along a branch in *T*

Genotypes may change along a branch *b* = (*u, v*) due to mutations occurred on *b*. We let the genotypes at *u* (respectively *v*) be (*g*_0_, *g*_1_) (respectively 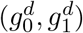). We use a *continuous-time Markov process* to model the process of (*g*_0_, *g*_1_) transiting into 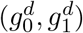
. The probability of such transition is called the *genotype transition probability* for any two genotypes. We now derive the transition probability recursively, using the *first-step analysis* based on the *memoryless* property of the exponential distribution,

There are two cases: (1) the transition probability 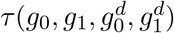 given that the branch has **no** PM, and (2) the transition probability 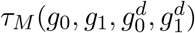 given that the branch **has** the single PM. In either case, CNM may occur. Due to the space limit, we only describe the case 1. 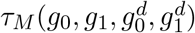 is derived in the Supplemental Materials (Sect. A.1).

##### Case 1: no PM along the branch

For a boolean variable *x*, we define an indicator function **I**(*x*) for *x*, where **I**(*x*) = 1 if *x* is true, and 0 otherwise. We define *γ*_*i*_(*g*_0_, *g*_1_) to be 1 if (*g*_0_, *g*_1_) allows a copy number *increase*, and 0 otherwise. Here, *γ*_*i*_(*g*_0_, *g*_1_) = 1 if *g*_0_ + *g*_1_ *< q*, and 0 otherwise. Similarly, we define *γ*_*d*_(*g*_0_, *g*_1_) to be 1 if (*g*_0_, *g*_1_) allows a copy number *decrease*, and otherwise. Here, *γ*_*d*_(*g*_0_, *g*_1_) = 1 if *g*_0_ + *g*_1_ *> p*. Recall that *p* and *q* are the lower and upper bounds of copy numbers, respectively. Then we have the following recurrences.

##### Case 1a

Either copy number increase or decrease is allowed: *γ*_*i*_(*g*_0_, *g*_1_) = 1 *or γ*_*d*_(*g*_0_, *g*_1_) = 1.

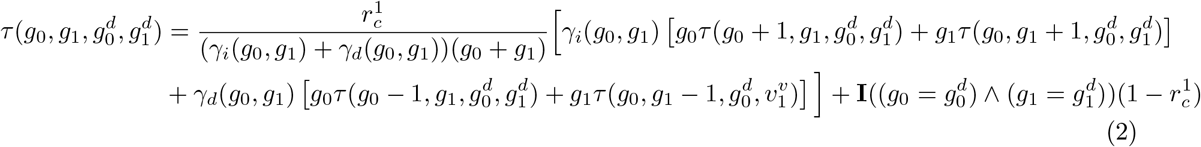

##### Case 1b

No copy number mutations is allowed: *γ*_*i*_(*g*_0_, *g*_1_) = *γ*_*d*_(*g*_0_, *g*_1_) = 0. Then, 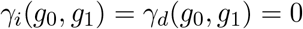 and 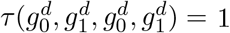 for all 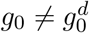 or 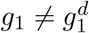.

##### Explanation

We focus on case 1a. We consider the *first* event forward in time. In case 1a, at least one copy number mutation is enabled. There are two possibilities for the first event. (i) The first event is a CNM. This occurs with probability *r*^1^ (Eq. 1). Recall that we assume that each CNM can only change (increase or decrease) the copy by one. Now suppose that it is an increase. Given that a CNM occurs, *regardless* whether both increase and decrease of copy number for the current genotype are allowed, the CNM increases copy number by one with probability 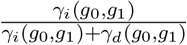. Furthermore, the CNM is an increase of 0 allele when the CNM picks one of the *g*_0_ 0 alleles among all *g*_0_ + *g*_1_ alleles. This happens with probability 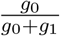 . Similarly, the CNM increases the number of allele with probability 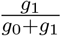. The case of copy number decrease is similar. (ii) The first event is a timeout event. This occurs with probability 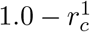 (Eq. 1). When a timeout occurs, if the source and destination genotypes are identical, the transition probability is 1, and 0 otherwise.

For example, Fig. 1(c) shows transitions of genotypes from the genotype (1, 1) along a branch in *T* within copy number range [1, 3]. Since the copy number of (1, 1) is 2, both copy increase or decrease are allowed: *γ*_*i*_(1, 1) = *γ*_*d*_(1, 1) = 1. Here, *g*_0_ = *g*_1_ = 1. Moreover, a CNM occurs as the first event with probability 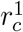 (Eq. 1). It may choose either the 0 or 1 allele to increase or decrease. Therefore, there are four genotypes, (0, 1), (1, 0), (2, 1) and (1, 2), that can be obtained by one CNM from (1, 1), each with the *same* probability 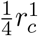. Moreover, a timeout can be the first (and terminal) event with probability 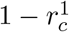 occurs and the genotype remains unchanged.

We calculate all possible genotype transition probabilities by solving the linear system formed by the above equations. Note that this calculation is performed *once* prior to the CLT inference. See Sect. A.2 in the Supplemental Materials for parameter estimation and other details.

## 3 Maximum likelihood inference of CLT on the JSC model

### 3.1 Data

We consider a set of *n* cells 𝒞 = {*c*_1_, *c*_2_, …, *c*_*n*_}, over a set of *m* SNV/CNA sites, 𝒮 = {*s*_1_, *s*_2_, …, *s*_*m*_} . The given data contains two parts: sequence reads ℛ and the estimated copy number profiles of the cells at these sites. The profile is represented by a matrix 𝒵 = {*z*_*i,j*_}, where *z*_*i,j*_ is the absolute copy number of the cell *c*_*i*_ at the site *s*_*j*_ and is assumed to be an integer. Note that for the same cell, different sites can have different copy numbers. For example, Fig. 1 part (e) shows data for *n* = 6 cells at *m* = 4 sites. For the cell *c*_3_ at site *s*_1_, there are two (respectively seven) reads for the wild-type (respectively mutant) allele, and the called copy number is two. That is, *z*_3,1_ = 2.

The data can be used to compute the *genotype likelihood L*(*G*; **Θ**) = *Pr*(ℛ, 𝒵|*G*, **Θ**) where the genotypes are constrained by the copy numbers 𝒵 for each cell at each site. Here, **Θ** is a set of parameters (e.g., allelic dropout rate) for scDNA sequencing. Intuitively, the higher a genotype likelihood is, the more likely this genotype is the true genotype for the site and cell. See Sect. A.3 in the Supplemental Materials on genotype likelihood calculation. For the rest of the paper, we assume that *L*(*G*; **Θ**) has been computed for all cells and sites.

### 3.2 Maximum likelihood inference

ScisTreeCNA aims to find the optimal CLT *T* ^*∗*^ and the best placement **b**^*∗*^ of PMs on *T* ^*∗*^ so that the likelihood *L*(*T*, **b**; ℛ, 𝒵, **Θ**) = *P* (ℛ, 𝒵 |*T*, **b**; **Θ**) is *maximized* (*i* is the site index):

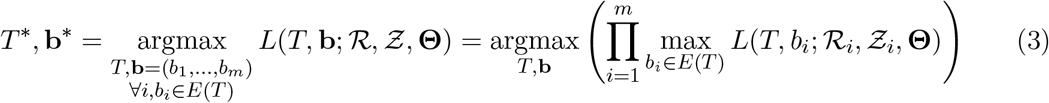

For each PM, we can place it on exactly one of the branches *b* ∈ *E*(*T*) (the set of edges in *T*). We define a sequence **b** = {*b*_1_, *b*_2_, …, *b*_*m*_} for the placement of PMs for all sites, where *b*_*i*_ ∈ *E*(*T*).

The key step is to compute the likelihood *L*(*T*, **b**; ℛ, 𝒵, **Θ**) of *T* and **b**. That is, we know on which branches of a given *T* PMs occur. In principle, *L*(*T*, **b**; ℛ, 𝒵, **Θ**) can be computed by *enumerating* all genotypes at *all* nodes of *T* ; for each set of enumerated genotypes, the likelihood of a site is equal to the product of all transition probabilities in *T* and the genotype likelihoods at leaves, i.e., extant cells, as well as the prior probabilities of the genotypes at the root of *T* . However, this would lead to an algorithm that runs in *O*(*nq*^4*n*^) time for a single site. This is clearly inefficient. See Sect.A.4 in the Supplemental Materials for details.

In the following, we present polynomial-time algorithms for evaluating and optimizing the likelihood. In the rest of this section, we focus on a single site due to the assumption of site independence. We drop the site indices to simplify the notations.

### 3.3 Algorithm for calculating the likelihood of a single site for a given *T*

We first present a dynamic programming-based algorithm with *O*(*nq*^4^) running time for computing *L*(*T, b*; ℛ, 𝒵, **Θ**) for a given tree *T* and the placement of the PM at a fixed site. On the high level, this algorithm resembles the well-known Felsenstein’s pruning algorithm [24]. We first give several definitions. Note that there is an *imaginary* branch entering the root of *T* . We define *L*_*v*_(*g*_0_, *g*_1_) for each valid genotype (*g*_0_, *g*_1_) to be the likelihood of observing (extant and ancestral) cells within *T*_*v*_ given that *v*’s genotype *G*_*v*_ is (*g*_0_, *g*_1_). We let *b* be the branch where the single PM at the focal site is located. When *v* is an internal node, we let *v*_*l*_ and *v*_*r*_ be the two children of *v*. There are three different cases about *v* and *b*:

**Case 1** *v* is a leaf in *T* . Here, the single extant cell’s genotype is fixed and its likelihood is *exactly* the genotype likelihood (Sect. A.3). So, *L*_*v*_(*g*_0_, *g*_1_) = *L*((*g*_0_, *g*_1_); **Θ**).

**Case 2** *v* is an internal node in *T, b* ≠ (*v, v*_*l*_) and *b* ≠ (*v, v*_*r*_) (i.e., the single PM is *not* on branches out of *v*, and so the genotype transition probability is *τ* instead of *τ*_*M*_).

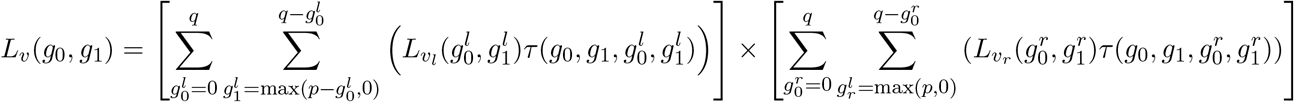

**Case 3** *v* is an internal node in *T*, and one of the branches out of *v* (say (*v, v*_*l*_)) is *b*. That is, the PM occurs along (*v, v*_*l*_) and so the genotype transition probability is *τ*_*M*_ instead of *τ* .

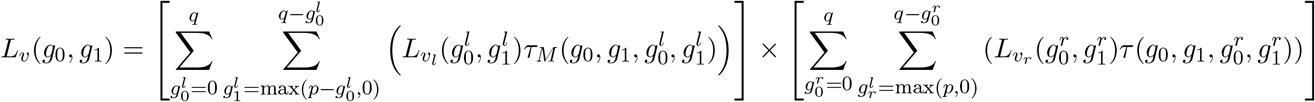

Then we obtain the likelihood of this site as *L*(*T, b*; ℛ, 𝒵, **Θ**) = *L*_root(*T*)_(*G*_root_).

#### Time analysis

There are *O*(*n*) subtrees in *T* . For each subtree, the size of the dynamic programming table *L*_*v*_(*g*_0_, *g*_1_) is *O*(*q*^2^). Calculating each of the values in this table takes *O*(*q*^2^) time. Therefore, assuming that the transition probabilities have been pre-computed, the likelihood at a single site can be calculated in *O*(*nq*^4^) time.

### 3.4 More efficient algorithm for optimally placing the mutations on a tree

The algorithm described in Section 3.3 assumes that the placement of the PM is given. Recall that, for a given tree *T*, we must also determine the branch *b*^*∗*^ on which to place the single PM that maximizes the likelihood (the inner optimization step in Eq. 3). Specifically,

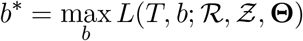

There are *O*(*n*) branches to place the single mutation. Naïvely, we would enumerate all choices of *b*. Therefore, the time for finding the maximum likelihood is *O*(*n*^2^*q*^4^) if we directly run the algorithm in Sect. 3.3 to find *b*^*∗*^. We now show that *b*^*∗*^ (and maximum likelihood) at a single site for a fixed tree *T* can be found in *O*(*nq*^4^) time, which is *one order of magnitude* faster than the above naïve approach.

#### High-level idea

The key observation for speeding up is that there are many *duplicate* computations for different choices of *b* if we calculate likelihoods from scratch. For example, suppose that the single PM occurs *outside* a subtree *T*_*v*_ (rooted at a node *v*) for two choices of *b*. Then when *v*’s genotype remains unchanged, the likelihood values are also unchanged within *T*_*v*_ for these two choices. Furthermore, the single PM occurring at *b* divides nodes of *T* into two parts: those inside *T*_*v*_ (i.e., IN) and those outside *T*_*v*_ (i.e., OUT). We can then apply dynamic programming again based on the placement of PM and the genotypes at the root of the subtrees.

#### IN and OUT

For each internal node *v* and one of its children *v*^*t*^, We have the following definitions: for genotype (*g*_0_, *g*_1_) where *g*_0_, *g*_1_ *≥* 0 and *p ≤ g*_0_ + *g*_1_ *≤ q*, and for a node *v* in *T*,

1. IN_*v*_(*g*_0_, *g*_1_):, the probability of the data (reads and copy numbers) of the cells *within T*_*v*_ given that *v*’s genotype is (*g*_0_, *g*_1_) and the single PM is *not inside T*_*v*_.
2. OUT_*v*_(*g*_0_, *g*_1_): the probability of data of all cells *outside T*_*v*_ given that *v*’s genotype is (*g*_0_, *g*_1_) and the single PM does *not* occur at any branches *outside T*_*v*_.

We have the following recurrences based on the same high-level logic as those in Section. 3.3. For a leaf *v*, IN_*v*_(*g*_0_, *g*_1_) = *L*((*g*_0_, *g*_1_); **Θ**). For an internal node *v*,

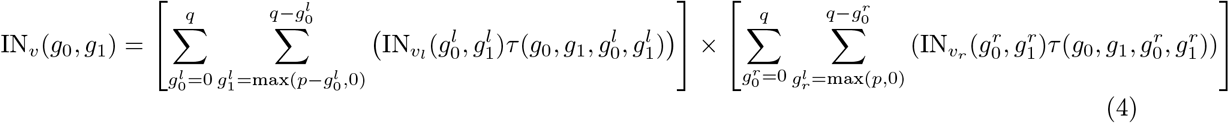

For the root(*T*) of *T*, OUT_root(*T*)_(*g*_0_, *g*_1_) = 1.0. For all other node *v*, let par(*v*) be the parent node of *v*, and sib(*v*) be the sibling of node *v*. Then,

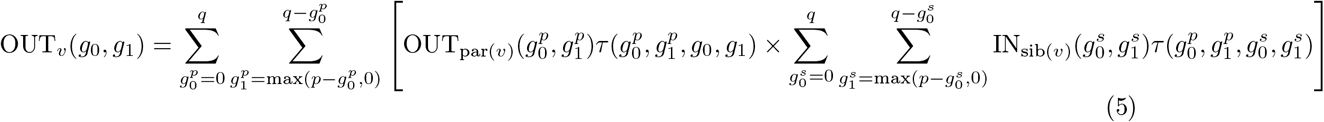

#### Find the best branch to place the single mutation

We first calculate all the values of IN_*v*_ and OUT_*v*_ using the above recurrences. We define 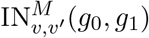 for each internal node *v* and one of its *children v*^*t*^, the probability of the data all the cells within *T*_*v*_ given that *v*’s genotype is (*g*_0_, *g*_1_) and the PM *occurs* along the branch (*v, v*^*t*^). The 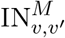 values are for the case when we know along which branch the PM occurs, and can be calculated from IN_*v*_ values by:

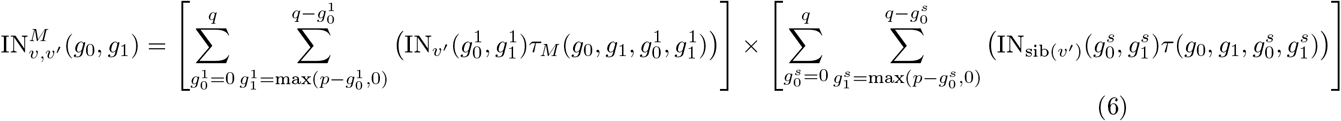

Then, for each choice of *b* = (*v, v*^*t*^), Eq. 7 shows that full likelihoods can be calculated by summing up the product of 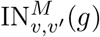 and OUT_*v*_(*g*) terms for all possible genotypes *g* at the node *v*. Thus, the best branch *b*^*∗*^ to place the single mutation at a site can be found by:

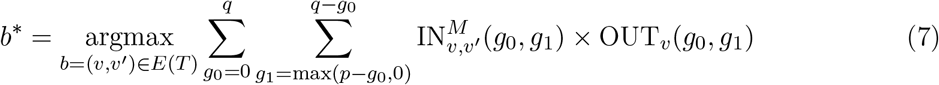

#### Time analysis

For a fixed site, calculating IN_*v*_ values takes *O*(*nq*^4^) time. 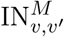 values can be calculated from IN_*v*_ in *O*(*nq*^4^) time. Directly calculating OUT_*v*_ from Eq. 5 takes *O*(*nq*^6^) time. However, the inner summations of Eq. 5 can be *pre-computed* for all possible genotypes 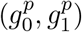 at *par*(*v*) *independently* from (*g*_0_, *g*_1_) in *O*(*nq*^4^) time. Therefore, OUT_*v*_ values can also be calculated in *O*(*nq*^4^) time. Therefore, finding *b*^*∗*^ and the maximum likelihood of a fixed site on a given tree can be done in *O*(*nq*^4^) time.

#### Local search

So far, we assume *T* is fixed. In practice, *T* is not known. To search for the optimal *T* ^*∗*^, we apply the local search, which starts from an initial *T* and moves to a neighboring tree with a higher likelihood at each iteration. See Sect.A.5 in Supplemental Materials for details.

### 3.5 GPU acceleration of local search

Finding the optimal *T* ^*∗*^ by local search using the algorithm in Sect. 3.4 can be further sped up using parallelism. The conventional parallelism uses multiple cores in CPU. ScisTreeCNA uses a *GPU* -based speedup approach. Note that most computation in searching for the optimal *T* ^*∗*^ is running the algorithm in Sect. 3.4 on the nodes of neighboring trees during local search. The *key* idea is that the algorithm in Sect. 3.4 can be converted into *matrix* operations, which can be performed efficiently by GPU using CUDA. Moreover, computations on two nodes in a tree can be processed *in parallel* in different CUDA cores if there is *no* ancestry relationship between them. The number of CUDA cores is usually in tens of thousands, while the number of CPU cores is often much smaller. Therefore, the GPU acceleration can be much more powerful than using CPU. See Sect. A.6 in the Supplemental Materials for details.

**Additional methods** are given in the Supplemental Materials due to the space limit.

#### Code availability

ScisTreeCNA is available at https://github.com/haotianzh/ScisTreeCNA.

## 4 Results

### 4.1 Simulated Data

We compared ScisTreeCNA with the following methods using simulated data: (i) ScisTree2 [7], CellPhy [6], (iii) DICE-star [11], and (iv) neighbor joining (NJ) [10]. ScisTree2 and CellPhy are the state-of-the-art SNV-based probabilistic inference methods. For methods that work solely with CNA data, we observed that neighbor joining based on Euclidean distance calculated from copy number profiles performs well in many cases. DICE-star is another distance-based method that was reported to outperform existing approaches using CNA data [11].

To evaluate performance and enable comparison, we used the following metrics (all normalized to the range [0,1], where a higher value indicates better performance):

- Tree Accuracy: The percentage of shared clades between the inferred trees and the true trees, which is equivalent to one minus the normalized Robinson–Foulds distance.
- Genotype Accuracy: The percentage of correctly called genotypes.

We first used CellCoal [25] to generate sequencing reads using its *infinite-site single-nucleotide deletions* feature. This can simulate data containing deletions only. The detailed script settings used for data generation with CellCoal are provided in Sect. B.1. See Sect. B.3 in Supplemental Materials about how the programs under comparison are run in simulation.

We simulated datasets with 50, 100, 150, and 200 cells. The coverage was fixed at 10, the dropout rate was set to 0.2, and the sequencing/amplification error rate was 0.01. In addition, we introduced a small amount of random copy number noise across all simulated datasets (see Sect.B.7 in Supplemental Materials). As shown in Figure S1 (in Supplemental Materials), ScisTreeCNA slightly outperforms the other four methods in both CLT reconstruction and genotype calling. However, CellCoal is limited to incorporating deletions only and therefore cannot fully reflect the performance of ScisTreeCNA, as copy gains also provide valuable information. To simulate data containing both copy number increases and decreases, we used a simulator called *scsim* that was originally developed in [5] and was released together with ScisTreeCNA. This simulator is logically similar to CellCoal (Supplemental Materials Sect. A.8 and Sect. B.2).

We evaluated all five methods on simulated data under the same settings described previously. Figure 2 shows that ScisTreeCNA significantly outperforms the other methods in tree accuracy when more copy number information is incorporated. This demonstrates that using both SNVs and CNAs can infer more accurate CLTs.

**Figure 2.**
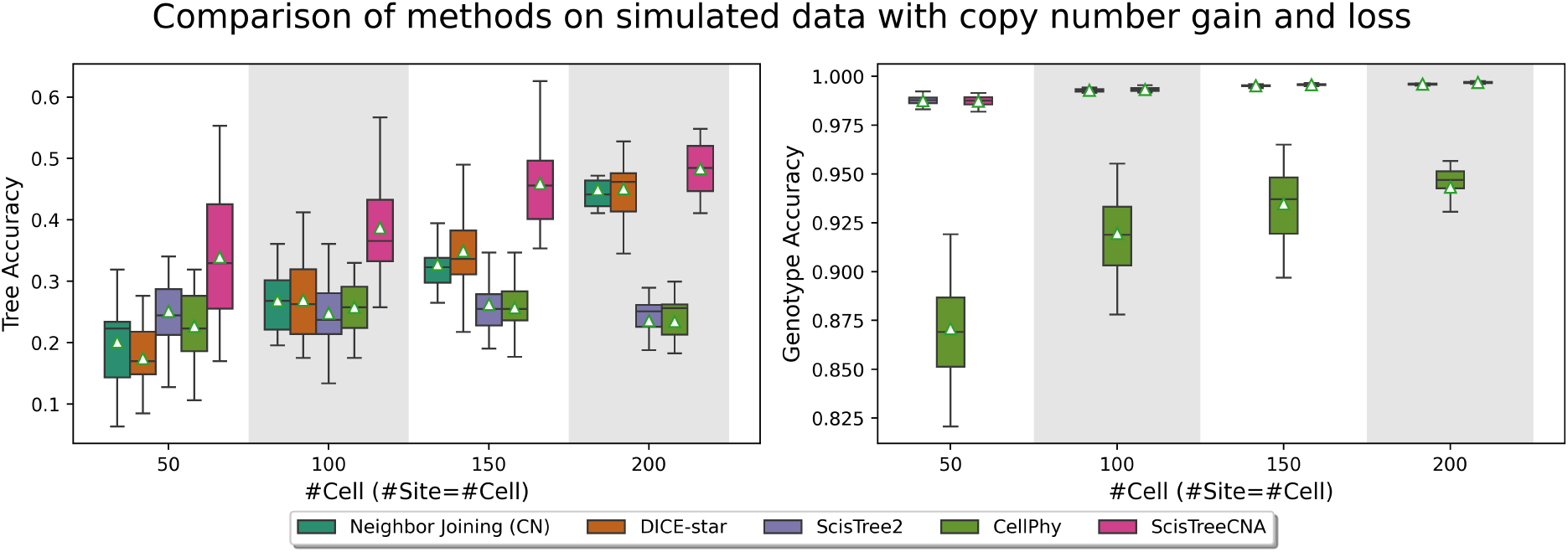
Boxplot of inference accuracy on data generated by scsim allowing both copy gain and loss. Number of cells: 50, 100, 150, or 200. Number of sites: equal to the number of cells. White triangle: the mean accuracy. Left: the tree accuracy. Right: the genotype accuracy (only ScisTree2, CellPhy and ScisTreeCNA call genotypes). Higher values indicate better performance in both.

#### Compare with ConDoR

So far, we only compared ScisTreeCNA with methods that use either SNVs or CNAs. There are existing cancer phylogeny inference methods, e.g„ ConDoR, that use both SNVs and CNAs and can produce CLTs given additional input such as the assignment of cells into clones. We compared with ConDoR using simulated data. The detailed results are shown in Sect. B.6. Overall, ScisTreeCNA outperforms ConDoR.

**More simulation results** are given in the Supplemental Materials due to the lack of space.

### 4.2 Real Data

#### 4.2.1 Data from Mission Bio’s Tapestri technology with three mixed cell lines

We evaluated ScisTreeCNA on an example dataset from Mission Bio generated using the Tapestri platform with a single-cell DNA AML panel [15]. The normalized percentages of sequencing reads across the panel amplicons were used to estimate CNAs for each gene analyzed. The dataset has 2, 476 cells profiled at 29 targeted loci, originating from three distinct cell lines: Jurkat, KG-1, and TOM-1. Jurkat cells were assumed to be pseudo-diploid over all genes and were used as a control line. The absolute ploidy of each site was then estimated based on this control. By mapping SNVs onto the CNA amplicons, we obtained the absolute copy number for each locus.

It is slow for ScisTreeCNA to analyze all 2, 476 cells. So we randomly subsampled 200 cells from the dataset, including 63 Jurkat, 82 KG-1, 35 TOM-1, and 20 cells labeled as “Mixed,” which are commonly regarded as doublets.

Figure 3a shows the inferred cell lineages reconstructed by ScisTreeCNA, ScisTree2, DICE-star and CellPhy. While the true CLT of these cells is not known, it is known that three cell lines are likely *not* closely related. Therefore, cells from each of the three cell lines are expected to *cluster* in the CLT. Both ScisTreeCNA and ScisTree2 show such clear clonal patterns in their inferred CLTs (near the roots), whereas CellPhy does not. Although CellPhy clusters cells from the same cell lines, its CLT suggests that TOM-1 diverged from Jurkat, which is *unlikely* because Jurkat is the control cell line. The CLT by DICE-star does not show clear clustering of cells from same cell lines, which is likely due to noise in estimated copy numbers.

**Figure 3.**
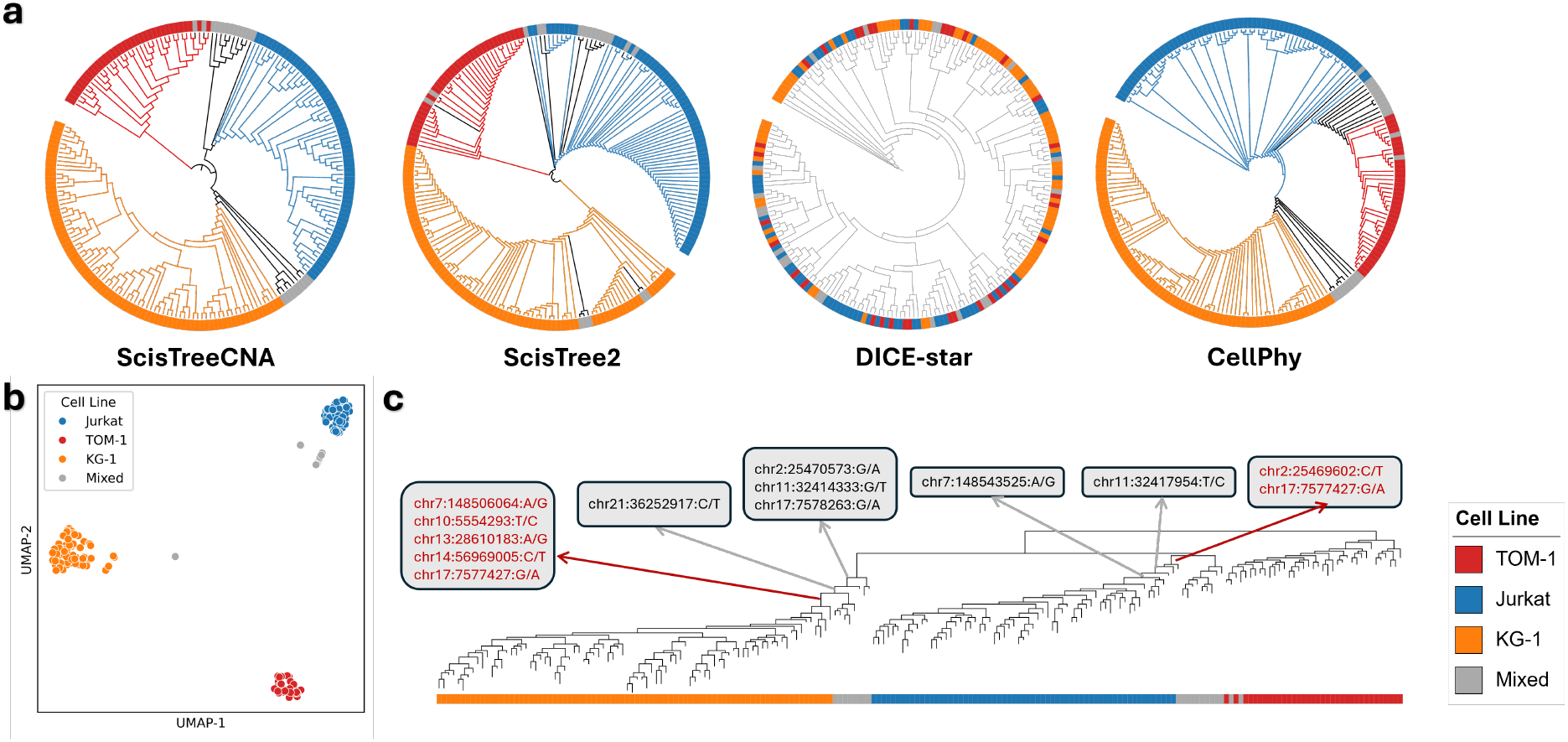
Comparison of inferred clonal lineage trees (CLTs) from ScisTreeCNA, ScisTree2, DICE-star, and CellPhy on the Mission Bio Tapestri dataset containing Jurkat, KG-1, and TOM-1 cell lines. Part (a): the reconstructed CLTs for ScisTreeCNA, ScisTree2, DICE-star, and CellPhy, with cell line indicated by color. Note that only the CLT by ScisTreeCNA has all three expected properties: (i) proper clustering of cells from a cell line, (ii) the expected absence of ancestral relations among the three cell lines and (iii) clustering of mixed cells. Part (b): UMAP clustering based on variant allele frequencies (VAFs), where cells labeled as “Mixed” represent putative doublets. Part (c): representative loss-of-heterozygosity (LOH) events mapped onto the ScisTreeCNA-inferred CLT. Genes highlighted in red denote key LOH variants that help distinguish doublets and infer proper clonal evolution among the three cell lines.

Figure 3b shows the clustering of cells using variant allele frequency (VAF). As indicated by the UMAP plot, mixed cells (possibly doublets) consist of mixtures of two cell lines that do not belong to any existing clusters. So mixed cells are expected to cluster in CLT. While the CLT inferred by ScisTreeCNA clearly shows the clustering of the mixed cells, these mixed cells do not cluster well in the CLT by ScisTree2. To further investigate how copy number influences tree construction, we mapped mutations exhibiting loss of heterozygosity (LOH) back to the ScisTreeCNA CLT (Figure 3c). This analysis reveals that cells from KG-1 share a set of LOH events (e.g., chr7:148506064:A/G) These LOH mutations may have helped ScisTreeCNA to infer more accurate CLT than ScisTree2 which does not use LOH information. Similarly, another group of LOH variants help to determine the relation between Jurkat and mixed cells.

#### 4.2.2 High-grade serous ovarian cancer single-cell DNA sequencing data

We ran ScisTreeCNA (and ScisTree2 and CellPhy) on a low-coverage targeted sequencing dataset derived from three clonally related high-grade serous ovarian cancer (HGSOC) cell lines from the same patient [26]. This data has 891 cells that were first aggregated into nine clones based on shared large-scale copy number alterations, and a clonal phylogeny was constructed using clone-specific SNVs (Figure 4a). We randomly sampled 200 cells and filtered out mutations missing in more than 75% of them, resulting in a dataset containing 200 cells and 721 mutations.

**Figure 4.**
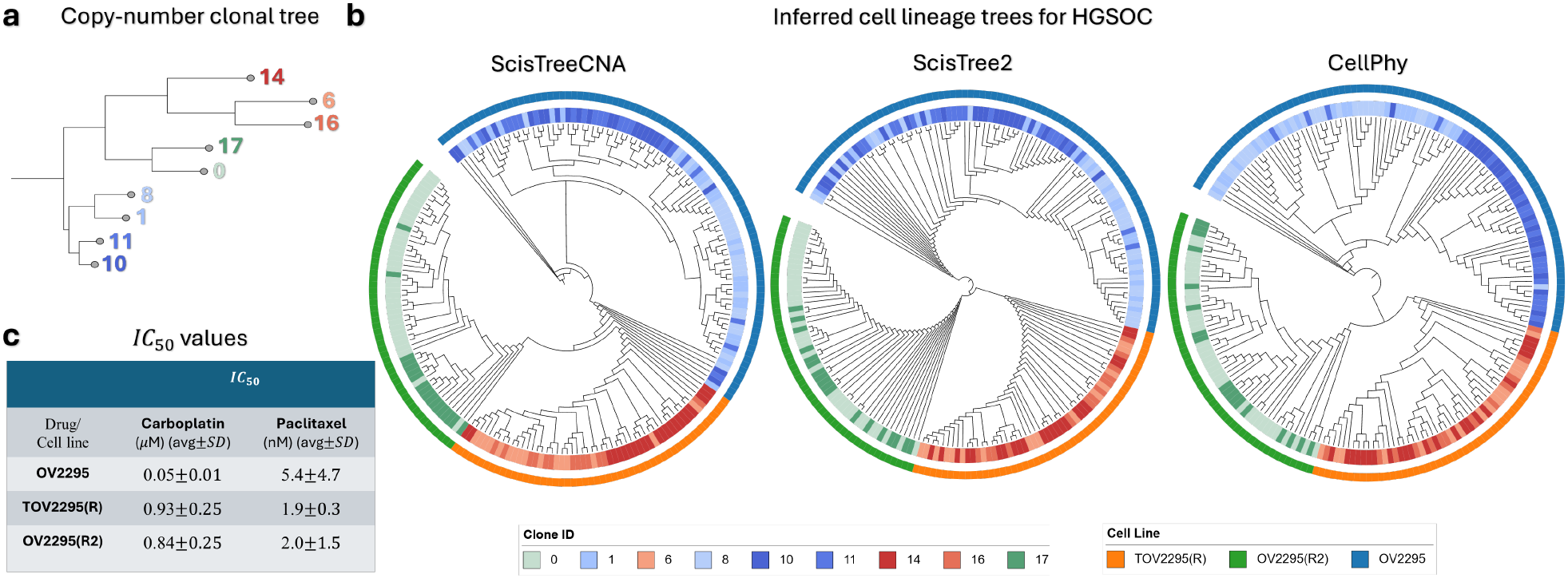
HGSOC data analysis. (a) Clonal phylogeny adapted from [26], in which nine clones were predefined based on copy number profiles. (b) CLTs reconstructed by ScisTreeCNA, ScisTree2, and CellPhy. The inner ring: copy-number-defined clones (shown in distinct colors). The outer ring: the three cell lines: OV2295 (primary ovarian tumor), TOV2295(R) (first recurrent tumor), and OV2295(R2) (second relapse tumor). The CLT inferred by ScisTreeCNA exhibits better clustering of cells belonging to the same clone compared to the other methods. Different from the other two CLTs, it also reveals that the second relapse, OV2295(R2), likely originated from the first relapse, TOV2295(R). (c) The IC_50_ values of each cell line for two drugs [27]. IC_50_ is the concentration of a drug that produces 50% of its maximum inhibitory effect: the higher value the greater drug resistance.

As shown in Figure 4(b), all three methods correctly grouped cells according to their cell line of origin. However, cells from the same assigned clone tend to be more dispersed in the CLTs inferred by ScisTree2 and CellPhy than that from ScisTreeCNA. For instance, cells from clones 14 and 17 cluster better in ScisTreeCNA’s CLT than in the other two CLTs.

We further investigated the evolutionary relationship among the three cell lines. OV2295 represents the primary ovarian tumor at initial diagnosis (pre-treatment), TOV2295(R) is the first recurrent tumor collected after chemotherapy, and OV2295(R2) is a second relapse tumor [27]. Both recurrent lines are known to originate from the primary tumor OV2295, which all three CLTs in Figure 4(b) are consistent with. However, the evolutionary ordering between TOV2295(R) and OV2295(R2) is less clear. As a second relapse, OV2295(R2) likely originated from a subclone within TOV2295(R), as both display similar drug resistance to Carboplatin and Paclitaxel and share comparable CNAs after chemotherapy, as shown in Figure 4(c) [27]. ScisTreeCNA’s CLT (but not those from ScisTree2 and CellPhy) supports a sequential progression model: TOV2295(R) split from OV2295, and then OV2295(R2) split from TOV2295(R).

## 5 Conclusions

Using SNVs alone for CLT inference may lead to low accuracy when some SNV mutants may be deleted due to copy loss and the IS model can be violated. Neglecting the copy gain information may miss true clades. Some existing scDNA datasets only have a small number of SNVs. Thus, using SNVs only may not infer a very refined CLT. On the other hand, using CNAs only may miss important clades revealed by SNVs. Moreover, noise in CNAs often reduces inference accuracy. Not using SNVs would miss the chance of using SNVs to offset the noise in CNAs. By combining SNVs and CNAs, ScisTreeCNA can infer a more accurate CLT than methods using only one type of variants. This is enabled by the JSC model, a novel probabilistic model for joint evolution of SNVs and CNAs, and related efficient algorithms for inference.

## Supplemental Materials

### A Supplemental methods

#### A.1 The case 2 of the evolution of genotypes along a single branch of CLT by the JSC model: the single PM occurs at the branch

In Sect. 2.2.2, we only derive the genotype transition probability for the case where the PM does *not* occur along a branch *b* in *T* . We now derive the case 2 where the PM does occur along *b*.

We denote the genotype transition probability along *b* where the single PM for the current site *s occurs* on *b* as 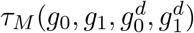. Since PM must occur on *b*, timeout is *not* allowed prior to the occurrence of the PM.

##### Case 2a *g*_1_ ≥ 1

Recall that the PM changes a 0 allele to a 1 allele. So in this case, a PM already occurs on a branch that is ancestral to *b*. Due to the assumption that the PM can occur only once, for all 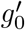 and 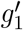,

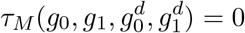

##### Case 2b

Either *γ*_*i*_(*g*_0_, 0) = 1 (copy number increase is allowed), or *γ*_*d*_(*g*_0_, 0) = 1 (copy number decrease is allowed). That is, at least one CNM is enabled. Note that this implies *g*_0_ ≥ 1. Also, *after* the PM occurs, due to the memoryless property of Markov process, we return to the Case 1 in the main text (Sect. 2.2.2). Then, by similar probabilistic reasoning as before,

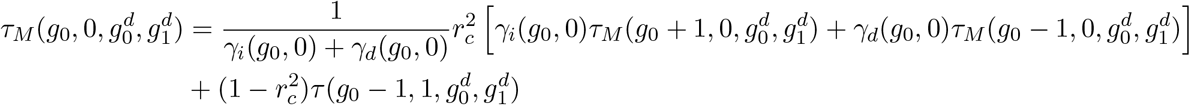

##### Case 3

*γ*_*i*_(*g*_0_, 0) = *γ*_*d*_(*g*_0_, 0) = 0. That is, no CNM is allowed for the genotype (*g*_0_, 0). Then the first event must be a PM with probability 1. So,

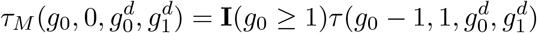

#### A.2 Solving Genotype Transition Distribution

By the JSC model, the genotypes form a two-dimensional *random walk* with an absorbing barrier (the timeout). The JSC model specifies the probabilistic rules in the form of a lienar system about how genotypes evolve along a branch of a CLT. To calculate the likelihood for a specific CLT *T*, we need to first calculate the transition probability of *all* possible genotypes for a single branch of a CLT.

##### Parameter estimation

We first estimate three parameters, namely Poisson rates: *λ*_*s*_ for point mutations, *λ*_*c*_ for copy number mutations and *λ*_*t*_ for timeout. Focusing on a single site, each branch in the tree *T* corresponds to one timeout event. Since *T* is a binary tree with *n* leaves, there are 2*n* − 1 branches (and hence timeouts) in total, so we let *λ*_*t*_ = 2*n* − 1. Under the IS model, only a single mutation occurs across the entire tree. Therefore, we simply set *λ*_*s*_ = 1. Estimating *λ*_*c*_, which controls the rate of copy number changes, is more nuanced. To estimate *λ*_*c*_, we apply the maximum parsimony (MP) method, a standard approach for reconstructing copy number trees using only the total copy number information of observed cells. Given a CLT *T*, we compute the *minimum* number (denoted as 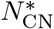) of total copy number changes for all *m* sites within *T* . Here, the copy number changes *N*_CN_ for fixed genotypes in *T* is calculated by *summing* up the absolute differences in the copy numbers at the source and the destination of all branches of *T* and all *m* sites. We let the copy numbers at leaves of *T* be the called copy numbers. 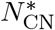 is the *minimum* of *N*_CN_ over all possible copy number settings at internal nodes of *T*, which can be calculated by standard dynamic programming for a fixed *T* in linear time. We then set 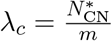.

##### Calculating transition probabilities

We consider two types of transition distributions: *τ*, representing transitions without PM, and *τ*_*M*_, representing transitions with PM. For each valid genotype, we formulate two corresponding equations: one under the PM model and one under the non-PM model. These equations can be organized into two separate *linear systems*. To obtain the actual transition distributions, we use the “scipy.linalg.solve” Python function to solve the linear systems and obtain the transition probabilities. The dimension of this linear system is *O*(*q*^4^). Note that we first solve for *τ*, followed by *τ*_*M*_, since the computation of *τ*_*M*_ depends on the probabilities obtained from *τ* .

#### A.3 The generalized genotype likelihood model

We calculate the likelihood of a given genotype for cell *i* at site *j*. To simplify the notations, we omit the indices of cells and sites in the following discussion with the understanding the reads and copies are for one specific site and cell. Specifically, we compute the likelihood of genotype *G* given the observed read counts *R* = (*r*_0_, *r*_1_), the called copy number *z*, and a set of parameters **Θ** including dropout rate, sequencing error rate, etc. Here, the observed read counts are *r*_0_ (reference) and *r*_1_ (mutant), the called copy number is *z*, and the true total copy number (unobserved) is 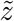. We impose bounds on the true copy number 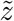, such that 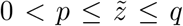. While sites with zero copies 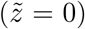 may occur in real data, we ignore this case in the current implementation.

Instead of working on conventional binary (0, 1) or trinary (0/0, 0/1, 1/1) genotype representations, we extend the genotype definition to a generalized genotype that explicitly specifies copy number. This generalized genotype, denoted by *G* = (*g*_0_, *g*_1_), is defined by the absolute copy numbers for the reference allele (*g*_0_) and the mutant allele (*g*_1_). Crucially, this definition is constrained such that the total number of copies equals the true copy number at that site: 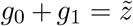. Note that all genotypes in this paper refer specifically to the generalized genotypes as defined above.

Then the genotype likelihood can be formally written as:

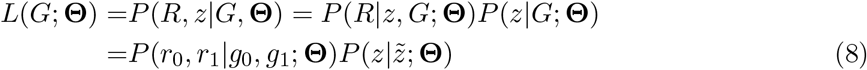

#### A.3.1 Copy number likelihood

We first consider the copy number likelihood 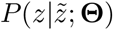. We model the deviation of the observed copy number *z* given the true copy number 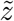 using a mixture of a degenerate distribution and a Poisson distribution. The purpose is allowing *noise* in the observed copy number. The copy number error rate is *ξ*, which is equal to the probability of copy numbers that are determined by a mixture of degenerate Poisson distributions. A special case is that the true copy number of zero 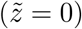 but the observed copy number is greater than zero (*z >* 0). This may occur due to noise. However, the structure of our mixture model does not permit the case where *z >* 0 *and* 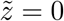 in this specific case. Consequently, we require 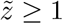.

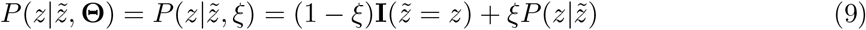

Here, **I** is the indicator function. 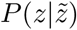 is the Poisson distribution with parameter 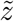.

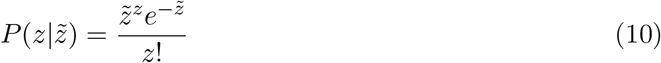

Plugging Eq. 10 back into Eq. 9, we obtain the copy number likelihood:

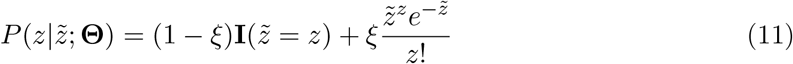

#### A.3.2 Reads likelihood

The reads likelihood *P* (*r*_0_, *r*_1_ | *g*_0_, *g*_1_; **Θ**) accounts for allelic dropout and amplification/sequencing errors. Not all genotypes are valid since we have constraints by copy numbers 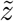. Thus, we define 𝒱 (*g*_0_, *g*_1_) for genotype (*g*_0_, *g*_1_):

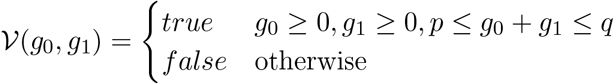

When 𝒱 (*g*_0_, *g*_1_) = *true*, we say that the genotype (*g*_0_, *g*_1_) is valid. We only consider valid genotypes throughout the paper.

##### Allelic dropout

We denote the dropout rate as *δ*. After dropout, 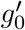 wild-type and 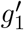 mutant alleles remain for sequencing. *g*_0_^*t*^ and *g*_1_^*t*^ are called the *effective allelic copy numbers*. This process is modeled independently for each allele using a Binomial distribution: *P* (*g*^*′*^ | *g*) ∼Binomial(*g*, 1 −*δ*).

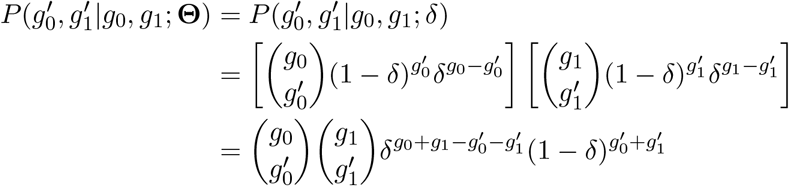

While it is possible to observe a very small number of reads even at a fully deleted site (i.e., when 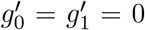) due to artifacts such as contamination or mapping errors, probabilistically modeling of reads in this specific scenario is non-trivial. We therefore require at least one amplified copy 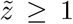 and neglect the case where 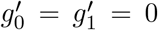. The probability of this full dropout is 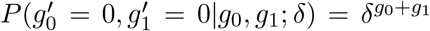. We normalize the probability distribution to condition on 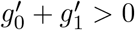:

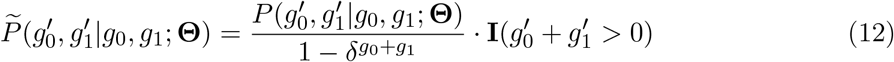

##### Sequencing error

We introduce a binary amplification/sequencing error model with error rate *ε*. Let *α ∈ {*0, 1*}* denote the true allele (reference=0, mutant=1).

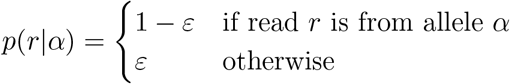

Given 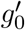 and 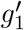, the likelihood of observing *r*_0_ reference and *r*_1_ mutant reads is:

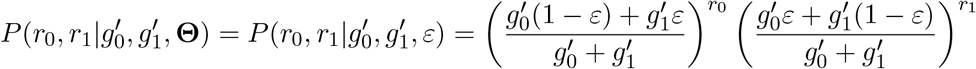

##### Total likelihood

Finally, we marginalize over all possible effective allelic numbers 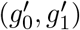 using the corrected probability 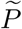 (Eq. 12):

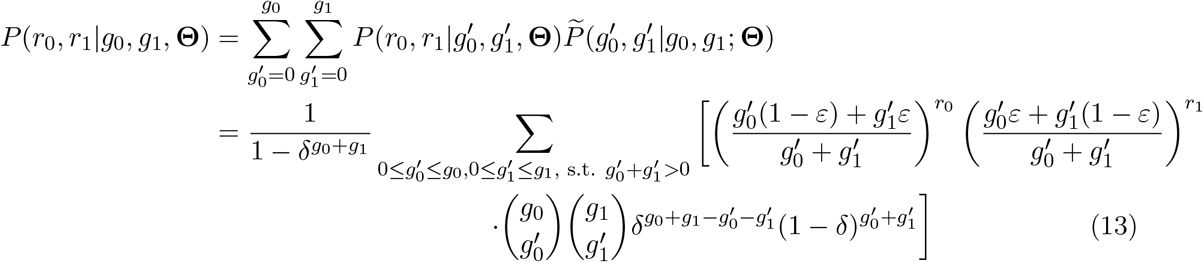

The full genotype likelihood now can be calculated by combining the copy number likelihood (Eq. 11) and the reads likelihood (Eq. 13) into Equation 8.

### A.4 Likelihood of JSC model for a given tree *T* at a specific site

To compute the likelihood of the JSC model *L*(*T, b*; ℛ, 𝒵, **Θ**) at a given site, let 𝒢 = *{G*_*v*_; *v ∈ V* (*T*)*}* denote the set of genotypes for all nodes in the tree *T* . For each node *v*, let *G*_*v*_ be its genotype. Let 𝒢 _leaf(*T*)_ denote the set of genotypes at the leaf nodes. By enumerating all possible genotype configurations 𝒢, we obtain

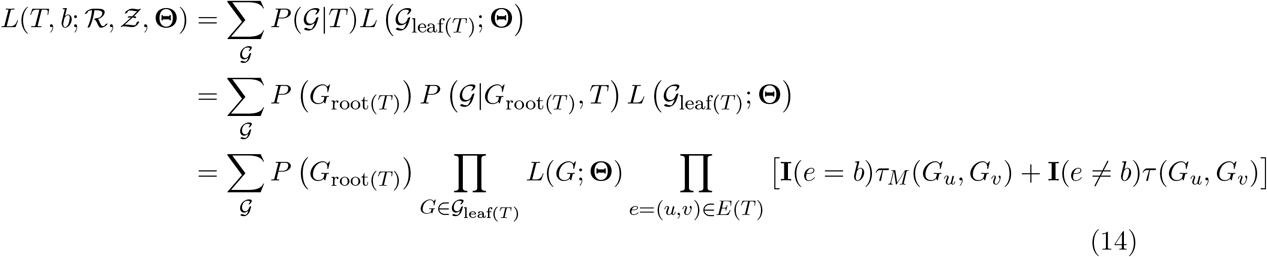

Here, **I** is the indicator function, *b* is the branch on which the PM occurs, and *E*(*T*) is the set of edges in *T* excluding the edge entering the root. We apply a non-informative prior to *G*_root(*T*)_. We illustrate the computation using the example in Figure 1(d). There is a 𝒢 annotation in the figure; for example, 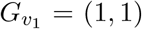 and 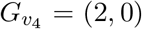. The branch on which the PM occurs is *b* = (root(*T*), *v*_1_), and the leaf genotypes are given by 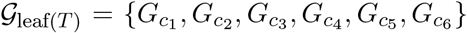. Then *P* (𝒢 | *T*) for this fixed assignment of 𝒢 can be computed as follows:

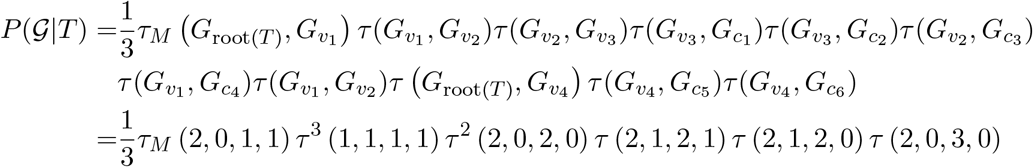

The genotype likelihood at leaves of *T* can be computed as follows,

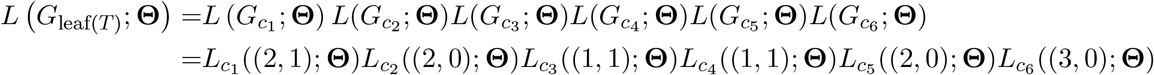

Here, the subscript indicates that the genotype likelihood is conditioned on the current cell. Evaluating Eq. 14 requires *O*(*n*) time for one specific 𝒢. For a given tree *T*, there are *O*(*q*^4*n*^) possible configurations of 𝒢 . Thus, the overall computational complexity for the total likelihood is *O*(*nq*^4*n*^).

### A.5 Local search

Local search is a widely used phylogenetic inference technique. At a high level, local search implemented in ScisTreeCNA works in a similar way as ScisTree and ScisTree2.

1. Construct an initial rooted binary tree *T*_0_ from the given data. Initialize *T*_opt_ *← T*_0_, *L*_opt_ *← L*(*T*_0_|ℛ, 𝒵).
2. Find rooted binary trees 𝒯_*c*_ that are within one tree arrangement operation (NNI or SPR) from *T*_opt_.
3. Let *T ∈* 𝒯_*c*_ that maximizes the likelihood *L*(*T* |ℛ, 𝒵). If *L*(*T* |ℛ, 𝒵) *> L*_opt_, set *T*_opt_ *← T, L*_opt_ *← L*(*T* |ℛ, 𝒵), and go to step 2. Otherwise, stop.

ScisTreeCNA uses the CLT reconstructed by ScisTree2 as the initial tree. ScisTreeCNA uses the NNI local search because NNI searches over a smaller tree neighborhood and allows more efficient local search than the SPR local search. We note that the JSC model is more complex than the IS model. Therefore, local search speedup tricks used in ScisTree2 are not easily applicable for ScisTreeCNA.

### A.6 GPU Acceleration

Computing the values 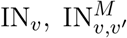, OUT_*v*_ for all *m* sites presents a significant computational bottleneck. As outlined in Section 3.4, this complexity is not only from the double summation loops over all possible genotypes, but also from the scale of the data, as the bottleneck grows linearly with the large number of sites (*m*). It is therefore natural to vectorize these calculations, transforming them into matrix operations that can be efficiently handled by modern computational frameworks such as OpenBLAS or Numba. Given the powerful matrix operation capabilities of GPUs, which currently dominate many scientific computing areas, vectorization allows us to fully leverage GPUs. By calculating the site-specific values in parallel during local search, this approach achieves a magnitude-level improvement in computational efficiency.

#### A.6.1 Vectorization

ℐ 𝒩 _*v*_ **and** ℐ 𝒩 _*v,v′*_ We consider all *m* sites and all valid genotypes simultaneously. Let 𝒢 = *{*𝒢_1_, 𝒢_2_, …, 𝒢_*k*_*}* denote the set of all *k* valid generalized genotypes.

For any given node *v* in the lineage tree, we define the matrix ℐ 𝒩 _*v*_, ℐ 𝒩 _*v,v′*_ ∈ ℝ^*m×k*^, where ℝ is the set of real numbers. The element (ℐ 𝒩_*v*_)_*ij*_ represents the value IN_*v*_(𝒢 _*j*_) at site *s*_*i*_ for node Similarly, the element (ℐ 𝒩 _v,v′_)_*ij*_ represents the value 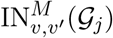 at site *s*_*i*_ for node *v* and one of its children *v* ^*′*^.

We define two *k × k* transition probability matrices:

1. Φ (no PM). The element Φ_*ij*_ denotes the probability of a genotypic transition from 𝒢 _*i*_ to 𝒢 _*j*_ along a branch where no PM occurs at that site.
2. Φ^*t*^ (with PM). The element 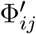 denotes the probability of a genotypic transition from 𝒢 _*i*_ to 𝒢 _*j*_ along a branch where a PM occurs at that site.

Then Eq. 4 is equivalent to

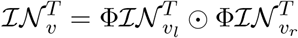

Eq. 6 is equivalent to

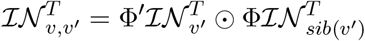

where ⊙ is the Hadamard product.

𝒪 𝒰 𝒯_*v*_ For the root of *T*, we define 𝒪 𝒰 𝒯_root(*T*)_ ∈ ℝ^*m×k×k*^, where each slice _root_[*i*] is the *k* × *k* identity matrix for *i* = 1, …, *m*.

For any *v* other than root (*T*), Eq. 5 is equivalent to

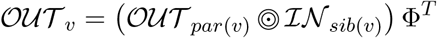

For a matrix 𝒪 𝒰 𝒯 ∈ ℝ^*m×k×k*^ and *IN* ∈ ℝ^*m×k*^, we define the operator 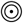 as batched row-wise Hadamard product, which produces a matrix of the same shape as 𝒪 𝒰 𝒯 .

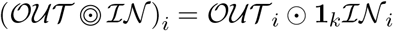

where *i* ∈ *{*1, 2, …, *m}*, **1**_*k*_ ∈ ℝ ^*k*^ is a column vector of ones.

##### Finding the best branches in batch

For any branch *b* = (*v, v* ^*′*^), we calculate:

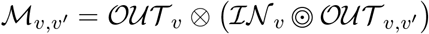

where ⊗ is batched matrix multiplication:

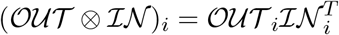

Then, following the same high-level logic as in Eq. 7, for any given site *s*_*i*_, the best placement *b*_*i*_ can be obtained by

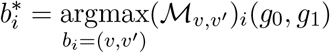

#### A.6.2 Batch Evaluation for Trees

Although computing ℐ𝒩, 𝒪𝒰𝒯 and ℳ values across multiple sites on a single tree using a GPU can be efficient, this approach does not fully utilize GPU resources. This limitation arises from the inherent dependencies within the tree structure. For example, computing ℐ 𝒩 of a node *v* requires prior knowledge of ℐ 𝒩 of its child nodes. These dependencies reduce the level of parallelism achievable when processing nodes within a single tree, limiting potential acceleration. To achieve speedup, evaluating multiple trees simultaneously enables more effective utilization of GPU. We developed an algorithm called Finding Independent Node Groups across Trees (FING) by first applying *topological sorting* to each tree as described in Algorithm 1, grouping nodes in a way that preserves their dependency constraints. An order parameter is used to control the traversal direction, either top-down (to calculate 𝒪 𝒰 𝒯) or bottom-up (to calculate ℐ 𝒩). We then merge independent groups across different trees to form larger, composite groups that can be processed in parallel by GPU. The pseudo-code of FING is given in Algorithm 2. Additionally, calculation of *ℳ* is not recursive and can be performed concurrently.

### A.7 Genotype Calling

Given the optimal tree *T* ^*∗*^ and the best mutation placement **b**^*∗*^, genotype calling is straight-forward. For each site, cells descending from a branch *b*^*∗*^ carrying the mutation are labeled as mutated (1).

It is also of interest to determine the most likely genotype assignments for all ancestral cells in *T* ^*∗*^. This can be achieved by finding the optimal 𝒢 ^*∗*^ at each site. This is similar to computing the likelihood in Sect.A.4. The difference is, instead of summing over all possible, we replace the summation with a maximization.

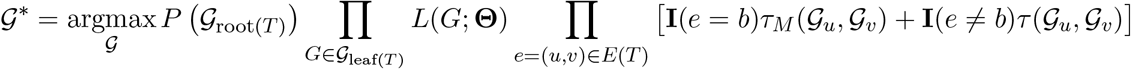

To compute this value, we use a dynamic programming algorithm, similar to the well-known Viterbi algorithm, which recursively finds the best genotypes for nodes in a bottom-up manner.

#### Algorithm 1

Topological Sorting for Finding Independent Node Groups in a Single Tree (TOPOSORT)

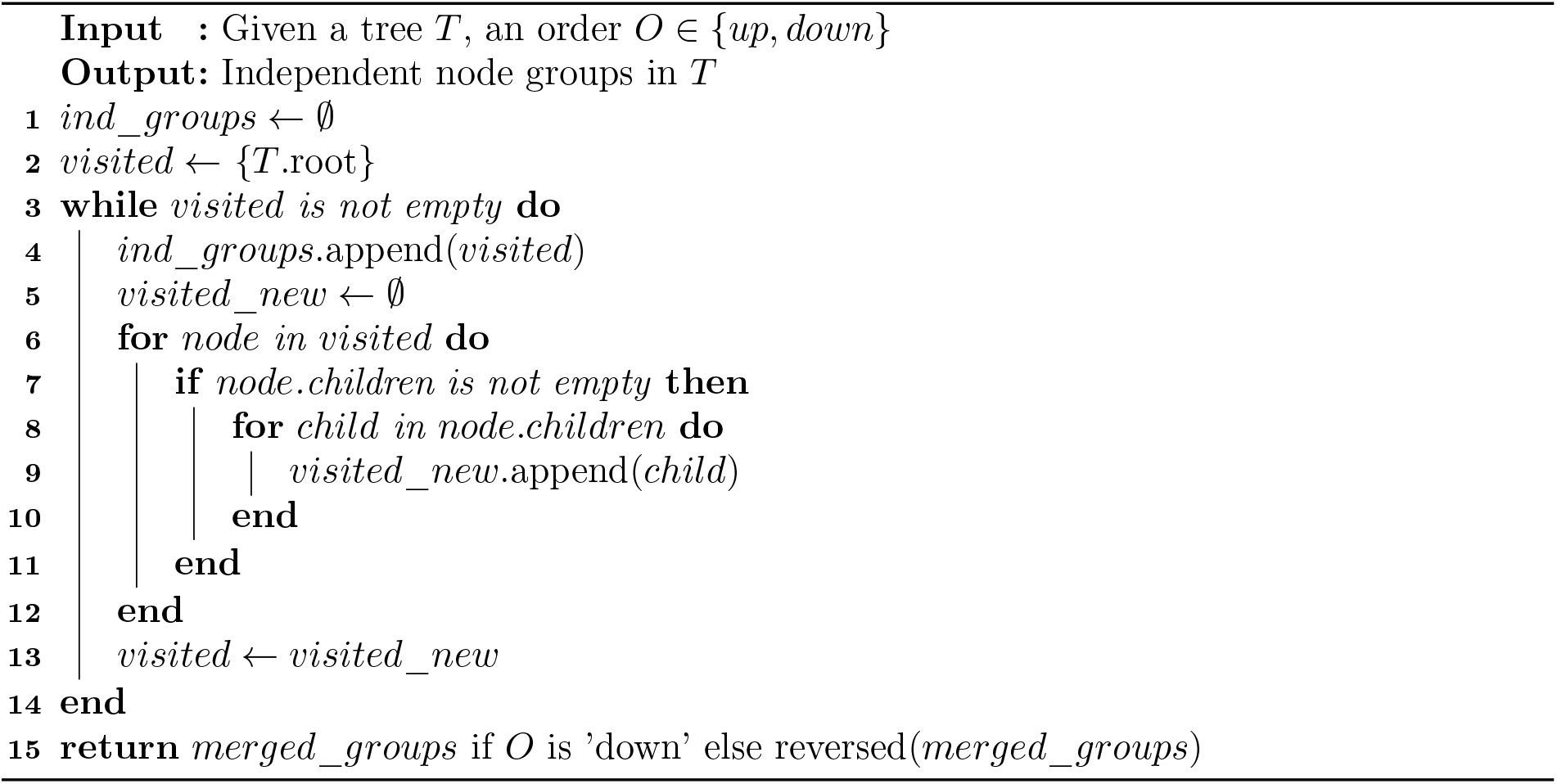

### A.8 Simulation of SNVs and CNAs with both copy gain and loss using scsim

In [5], we developed a simulator called *scsim* that can simulate SNV data with copy number changes. We have made small changes in its implementation. On the high level, it works as follows.

1. Given a fixed *T*, initialize the genotype at the root of *T* to be (2, 0) (diploid homozygous wild-type).
2. From the root, along each branch of *T*, simulate the PMs and CNMs using the standard Poisson processes (whose rates are given by the user) forward in time. Here, PM is only allowed once (the IS assumption). A CNM chooses an arbitrary allele and either makes a copy or deletes the allele with the same probability. Multiple mutations are allowed along a branch.
3. Once genotypes are generated at all the leaves of *T*, sequence reads are simulated. For each allele in the genotype of an extant cell, simulate allelic dropout by discarding the allele with a fixed dropout rate (i.e., no reads are generated if the allele is dropped out). If the allele is not dropped out, simulate reads with a small chance of sequencing error.

The code and detailed instructions are available at https://github.com/haotianzh/scsim.

## B Additional Results

### B.1 Simulation details of CellCoal

We follow the default simulation parameters described in the CellCoal manual (v1.1). Specifically, the effective population size was set to 10,000, the dropout rate to 0.2, with no doublets, a sequencing error rate of 0.01, and a sequencing coverage of 10. The deletion rate was set to 2.5×10^*−*6^, as recommended in Section 5.2.5 of the CellCoal manual. By default, CellCoal does not output copy number information. To obtain these data, we enable verbose mode by setting `-y3`, which lets CellCoal print calculation status and event logs. This allows us to trace deletion events along specific lineages and thereby obtain the corresponding copy numbers.

#### Algorithm 2

Algorithm for Finding Independent Node Groups across Trees (FING)

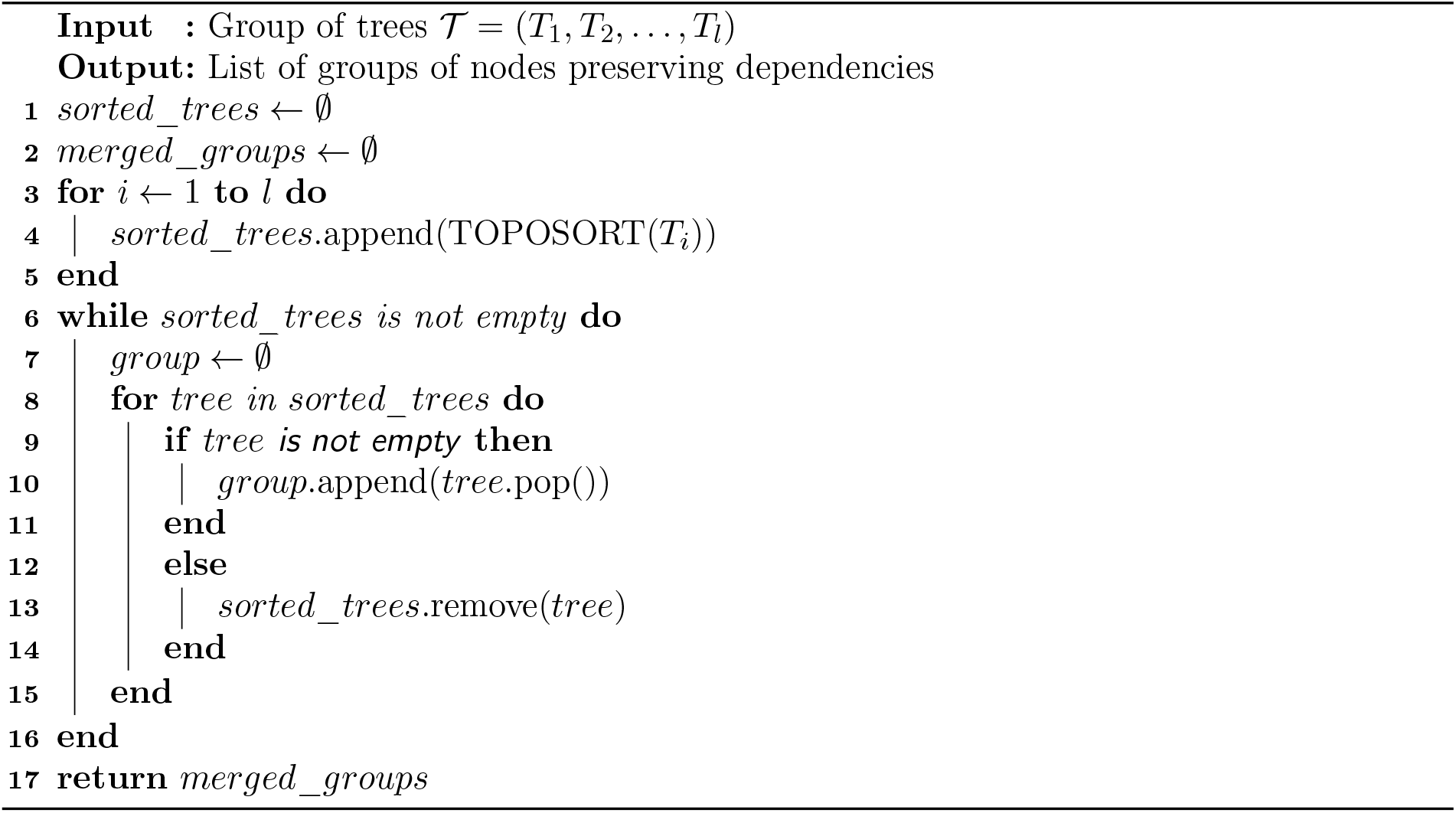

The example above simulates 20 replicates with 100 cells and 100 sites.

**Script 1.**
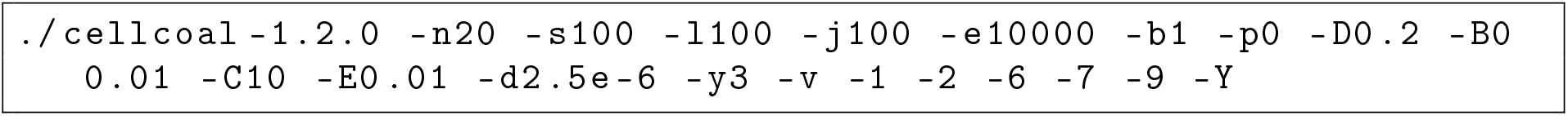
Example CellCoal simulation script.

### B.2 Simulation details of scsim

We followed the simulation protocol described in the `scsim` repository (https://github.com/haotianzh/scsim). By default, the sequencing error rate was set to 0.01, the dropout rate to 0.2, the copy gain rate to 0.05, and the copy loss rate to 0.03. Sequencing coverage was set to 10 with a dispersion of 5, and read counts were generated from a normal distribution.

This example simulates a dataset based on a tree with 100 leaves and 100 sites.

**Script 2.**
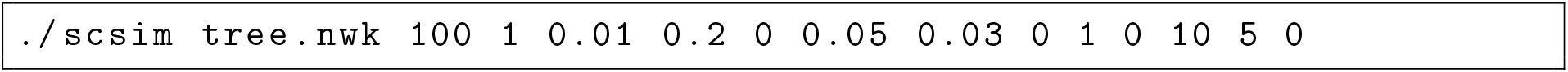
Example scsim simulation script.

### B.3 Running various programs in simulation

ScisTree2, CellPhy and DICE-star were all executed under their recommended optimal settings. Specifically, ScisTree2 was run with the SPR local search enabled; CellPhy is run in the “SEARCH” mode using the genotype likelihood (GT10) model; DICE-star was run with *DICE*^*∗*^ and the total copy number as input, following the data preparations and configurations described in their respective papers. The genotype likelihood for CellPhy was calculated in the same way as in CellCoal [25]. Moreover, since DICE-star and *NJ* are based solely on copy number data without incorporating SNV information, they do not perform genotype calling. All experiments were run on a LambdaLab Linux workstation with an NVIDIA RTX 4090 (24 GB VRAM), 32 CPUs, and 256 GB RAM.

### B.4 Simulated data by CellCoal with deletions only

Figure S1 shows the results on data simulated from CellCoal with deletions (no copy gains). As shown in Figure S1, when only deletions are present, ScisTreeCNA exhibits better performance than the other methods.

**Figure S1.**
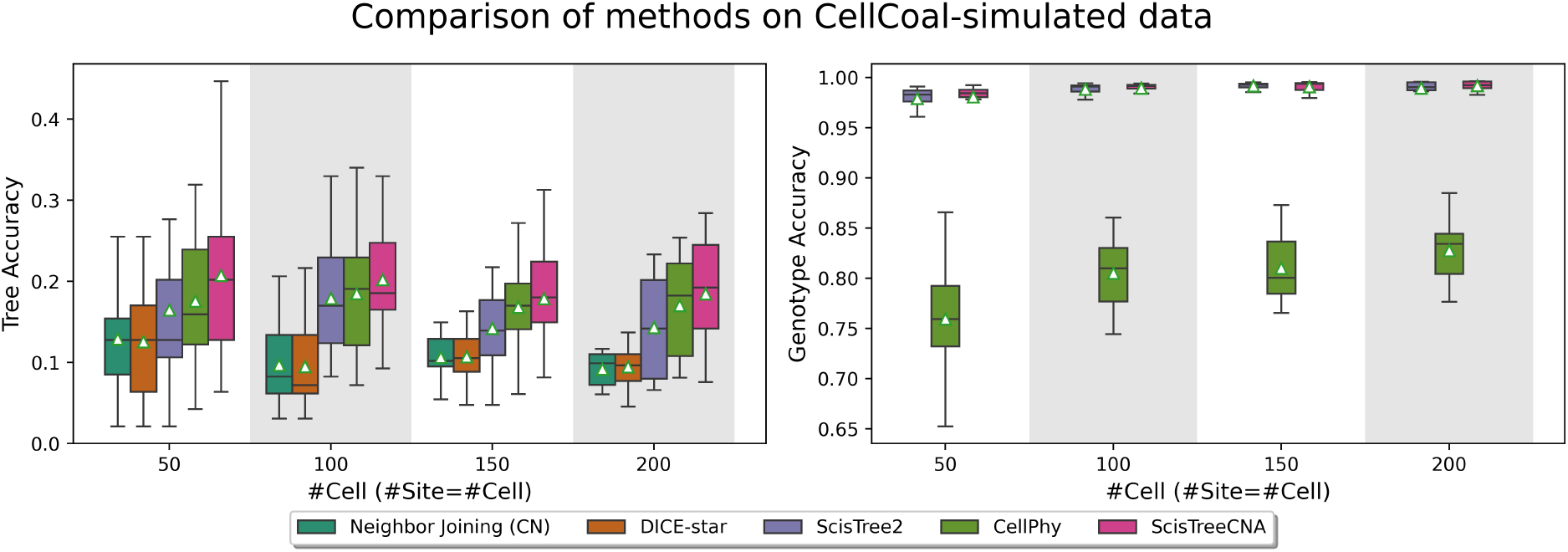
Boxplots of accuracy on datasets generated by CellCoal including only copy losses (no copy gains). Each dataset (20 replicates) contains 50, 100, 150, or 200 cells, with the number of sites equal to the number of cells. The white triangle indicates the mean accuracy. Left: tree reconstruction accuracy. Right: genotype accuracy (only ScisTree2, CellPhy, and ScisTreeCNA perform genotype calls). Higher values correspond to better performance in both metrics.

### B.5 Simulated data: comparison of data with clone-specific copy numbers

In practice, obtaining cell-specific copy number information is challenging due to limitations in sequencing technologies. As a result, researchers often pre-cluster cells into clones based on features of interest, such as known biomarkers, and derive clone-specific copy number profiles by averaging the values within each clone. To evaluate the performance of ScisTreeCNA for this scenario, we generated data with an underlying clonal structure, where cells within the same clone share an *identical* copy number profile. That is, the copy numbers are the average of cells in a clone and not for individual cells. For each combination of clone number and cell number, we simulated three conditions: 50, 100, and 150 cells for 4, 6, and 8 clones, respectively, with 10 replicates for each setting of parameters.

The results in Figure S2 show that even with clone-specific copy number profiles, ScisTreeCNA outperforms other methods. This suggests that clone-specific information plays a crucial role in correcting the topology at the clonal level.

**Figure S2.**
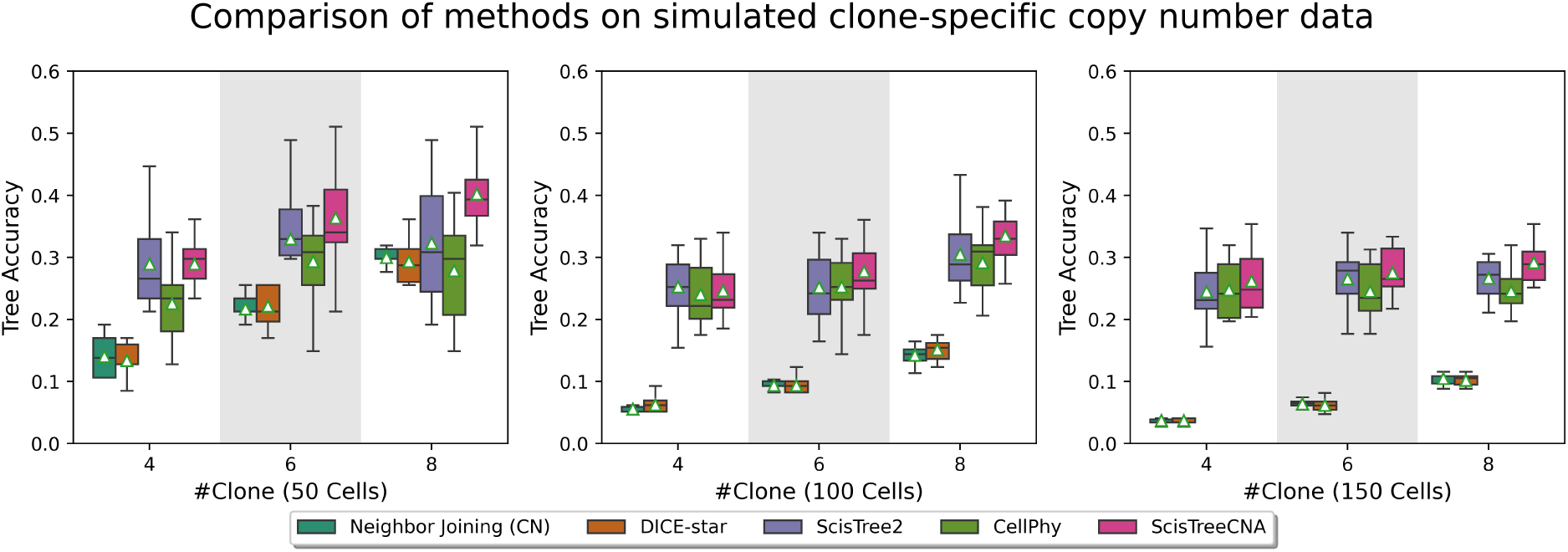
The boxplots for the tree accuracy of different methods for data simulated under a clonal tree structure. Varying numbers of clones and cells. The x-axis: the number of clones (4, 6, and 8 clones) across three cell counts (50, 100, and 150 cells), with 10 replicates per condition.

### B.6 Simulated data: comparison with ConDoR

So far, we have only compared ScisTreeCNA with methods using only one type of variants (SNVs or CNAs). Some existing tumor phylogeny methods use both SNVs and CNAs and can be adapted to produce CLTs when provided with additional input, such as a predefined copy-number tree or cell–clone assignments. In this study, we compared CLT inference methods with ConDoR[18], a state-of-the-art tumor phylogeny approach. In addition to read count data, ConDoR requires cell–clone assignments. We therefore supply the true assignments generated by scsim.

We encountered several challenges when running ConDoR: (i) convergence is not always guaranteed and can be extremely slow; (ii) it cannot handle a large number of clones; and (iii) as a *k*-Dollo model, it becomes inefficient when *k* is large (e.g., *k* = 5). To avoid non-terminating runs, we impose a 30-minute Gurobi time limit (the ILP solver used by ConDoR) and return the best solution found upon timeout. Because ConDoR outputs both a cell lineage tree and an imputed genotype matrix, we compare both tree and genotype accuracy in our comparisons.

We simulated data using scsim following the setup of Sect. B.5. However, we used fewer cells and clones: 25, 50, and 100 cells, each with 4 or 6 clones. For each configuration, we generate five replicates.

As shown in Figure S3, ConDoR generally achieves tree accuracy comparable to or slightly better than copy-number–based methods, which is consistent with its design as a clonal tree inference tool. It also tends to produce a more accurate imputed genotype matrix than CellPhy. Overall, ScisTreeCNA achieves the best performance across all metrics among the methods being compared.

**Figure S3.**
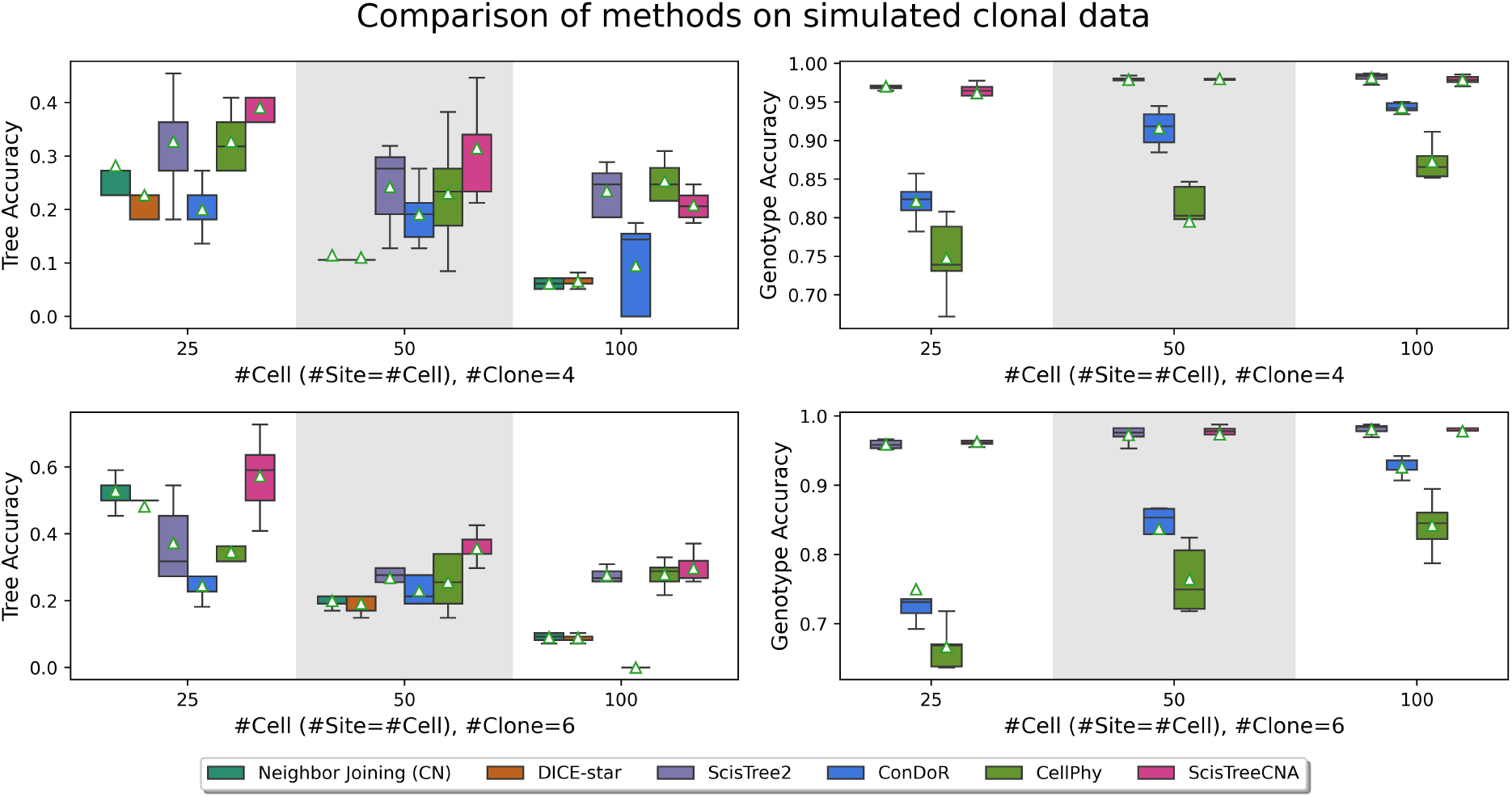
Comparison of CLT inference methods including ConDoR (a cancer phylogeny method) on simulated clonal data. Boxplots show tree accuracy (left) and genotype accuracy (right) for datasets simulated with 4 clones (top row) and 6 clones (bottom row). We vary the number of cells (25, 50, and 100), with the number of sites set equal to the number of cells. ConDoR is provided with true cell–clone assignments.

### B.7 Simulated data: robustness on copy number noise

To evaluate the performance of ScisTreeCNA under varying levels of copy number noise, we simulated datasets with noise levels of 0%, 10%, 20%, and 30%. Specifically, for each genomic locus, the true copy number had a corresponding probability (equal to the noise level) of being increased or decreased by 1 to introduce noise. In all experiments, the copy number error parameter of ScisTreeCNA was set to 0.05, without explicitly providing the true noise level for inference.

As shown in Figure S4, the accuracy of ScisTreeCNA and other copy-number-based methods drops with increasing noise levels. ScisTreeCNA consistently outperforms competing approaches for most cases noise but shows reduced accuracy at 30% noise, where copy number signals become severely distorted. We note that copy numbers in real data are believed to be more reasonably accurate.

**Figure S4.**
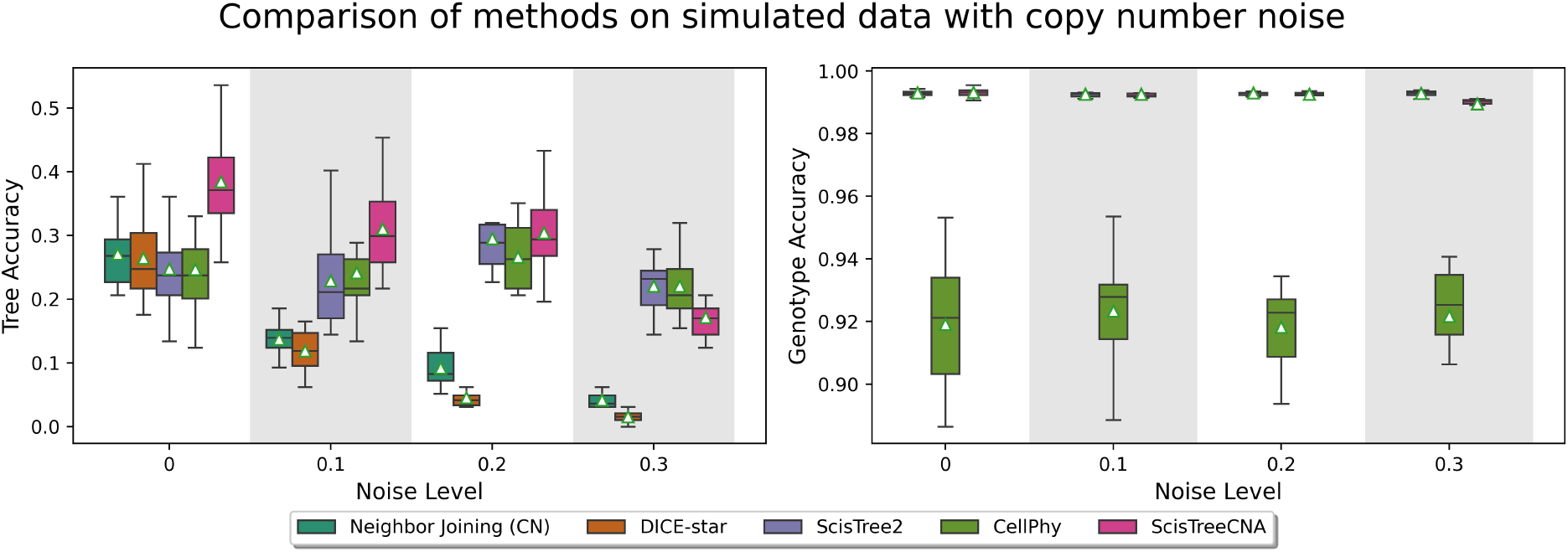
Accuracy comparison of ScisTreeCNA and other methods under varying levels of copy number noise. x-axis: copy number noise level between 0 and 30%. ScisTreeCNA consistently outperforms other methods at low to moderate noise levels, demonstrating robustness to copy number perturbations.

### B.8 Simulated data: robustness on incomplete copy numbers

We evaluated ScisTreeCNA under the scenario where copy number information is incomplete. In real data, copy numbers may be missing for some sites and cells. We simulated datasets with 100 cells and 100 sites, applying copy number masking rates of 10%, 30%, 50%, and 70%. For ScisTreeCNA, if a copy number is missing for any given site and cell, the copy number likelihood term described in Section A.3.1 is omitted, allowing the JSC model to transition flexibly among all possible genotypes. That is, when a copy number is missing, all copy numbers within the valid range are allowed.

As shown in Figure S5, the accuracy of ScisTreeCNA decreases as the proportion of missing copy numbers increases, due to the reduced availability of copy number information. Nevertheless, even under extremely high missing rates (e.g., 70%), ScisTreeCNA performs comparably to or better than ScisTree2, demonstrating robustness to incomplete copy number data. Moreover, compared with copy-number-only methods, the CLT inferred by ScisTreeCNA retains SNV information when copy number data are largely missing, yielding greater stability than neighbor joining and DICE-star which only use CNAs.

**Figure S5.**
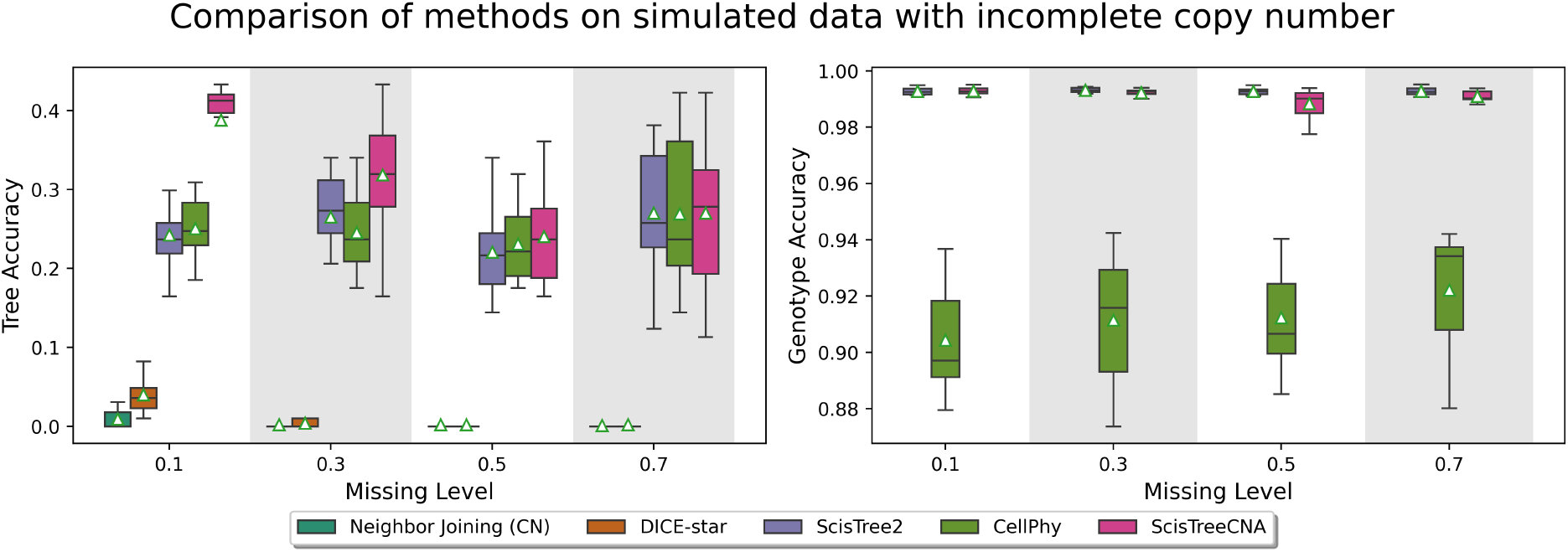
Accuracy comparison of ScisTreeCNA and other methods on simulated data with varying levels of missing copy number information. x-axis: copy number missing level between 10% to 70%. Left: tree accuracy. Right: genotype accuracy.

### B.9 Running time

In this section, we compare the running times of all methods on datasets of varying sizes. We tested datasets containing 50, 100, and 200 cells, each with 50, 200, and 500 sites. Neighbor joining and DICE-star were executed in single-thread mode, as they do not support parallelization. ScisTree2 and CellPhy were run using 30 threads, while ScisTreeCNA was executed on a single CPU thread with an RTX4090. As shown in Table 1, although ScisTreeCNA is the most time-consuming method due to its more complex models and algorithms, the runtime for moderate-sized datasets (e.g., 200 cells) remains acceptable. Notably, by leveraging GPU parallelization, ScisTreeCNA does not scale linearly with the number of sites. In fact, increasing the number of sites can even reduce runtime, as the additional information from SNVs and CNAs may accelerate model convergence.

**Table 1.** Running time (in seconds) of all methods on datasets of different sizes. Number of cells: 50, 100, and 200. Number of sites: 50, 200, and 500. Experiments were conducted on a LambdaLab workstation equipped with 256 GB RAM, 32 CPU cores, and an NVIDIA RTX 4090 GPU with 24 GB VRAM.

|  | CellPhy |  |  | Neighbor joining |  |  | DICE-star |  |  | ScisTree2 |  |  | ScisTreeCNA |  |  |
| --- | --- | --- | --- | --- | --- | --- | --- | --- | --- | --- | --- | --- | --- | --- | --- |
| #site | 50 | 200 | 500 | 50 | 200 | 500 | 50 | 200 | 500 | 50 | 200 | 500 | 50 | 200 | 500 |
| #cell |  |  |  |  |  |  |  |  |  |  |  |  |  |  |  |
| 50 | 3.21<br>($\pm 0.1$) | 3.48<br>($\pm 0.2$) | 3.92<br>($\pm 0.1$) | 0.05<br>($\pm 0.0$) | 0.06<br>($\pm 0.0$) | 0.06<br>($\pm 0.0$) | 0.25<br>($\pm 0.0$) | 0.25<br>($\pm 0.0$) | 0.28<br>($\pm 0.0$) | 0.04<br>($\pm 0.0$) | 0.08<br>($\pm 0.0$) | 0.12<br>($\pm 0.0$) | 40.25<br>($\pm 8.3$) | 15.52<br>($\pm 1.5$) | 8.17<br>($\pm 1.3$) |
| 100 | 6.91<br>($\pm 0.7$) | 7.70<br>($\pm 0.5$) | 8.36<br>($\pm 1.2$) | 0.39<br>($\pm 0.0$) | 0.39<br>($\pm 0.0$) | 0.39<br>($\pm 0.0$) | 0.30<br>($\pm 0.0$) | 0.30<br>($\pm 0.0$) | 0.35<br>($\pm 0.0$) | 0.09<br>($\pm 0.0$) | 0.17<br>($\pm 0.0$) | 0.37<br>($\pm 0.1$) | 558.58<br>($\pm 104.0$) | 223.75<br>($\pm 23.2$) | 103.87<br>($\pm 15.0$) |
| 200 | 17.89<br>($\pm 2.0$) | 13.64<br>($\pm 0.5$) | 15.45<br>($\pm 0.7$) | 1.27<br>($\pm 0.0$) | 1.27<br>($\pm 0.0$) | 1.29<br>($\pm 0.0$) | 0.38<br>($\pm 0.0$) | 0.39<br>($\pm 0.0$) | 0.41<br>($\pm 0.0$) | 0.13<br>($\pm 0.0$) | 0.29<br>($\pm 0.0$) | 0.60<br>($\pm 0.1$) | 1949.20<br>($\pm 551.7$) | 1242.48<br>($\pm 264.6$) | 490.38<br>($\pm 50.1$) |

